# Joint multi-omics profiling of brain and body health in aging

**DOI:** 10.1101/2025.10.09.681483

**Authors:** Asa Farahani, Zhen-Qi Liu, Filip Morys, Roqaie Moqadam, Yashar Zeighami, Mahsa Dadar, Alain Dagher, Bratislav Misic

**Affiliations:** Montréal Neurological Institute, McGill University, Montréal, QC, Canada; Department of Medicine, University of Montréal, Montréal, QC, Canada; Douglas Mental Health University Institute, Montréal, QC, Canada

## Abstract

The human brain and peripheral systems undergo coordinated changes throughout the lifespan, yet studies of aging have traditionally examined these systems as separate entities. Here we ask how brain health relates to peripheral biomarkers of bodily health including body mass index, blood pressure, and blood biochemistry results. We use partial least squares analysis to identify generalizable patterns of covariance between multi-modal neuroimaging data (structural, functional, diffusion, and arterial spin labeling MRI), demographic, and peripheral physiological markers in two large-scale deeply phenotyped datasets: the Human Connectome Project–Aging and UK Biobank. This data-driven pattern learning approach identifies two principal axes of brain–body associations in both biological sex groups. The first axis is driven by the dominant contribution of age. Across multiple brain measures, aging is associated with loss of brain structural integrity and cerebral vascular dysfunction. The second axis is driven by metabolic features, characterized by low high-density lipoprotein cholesterol, elevated body mass index, blood pressure, glycosylated hemoglobin, insulin, glucose, and alanine aminotransferase, and reduced cerebral blood perfusion. Finally, we show that deviations from a healthy metabolic profile are linked to cognitive deficits, particularly in females. Our study contributes to development of comprehensive translatable biomarkers for brain health assessment, and highlights the importance of metabolic health as a determinant of brain health.

## INTRODUCTION

The human brain and body are highly interconnected throughout the lifespan [7, 209]. Yet, the study of aging has typically treated cerebral and peripheral systems as separate entities. Currently, population-based aging trajectories are established for cortical thickness and gray matter density [27, 48, 60, 179, 195], brain volume and morphology [71, 90, 115], myelination [16, 76], structural connectome [49, 232, 245], functional connectome [19, 20, 32, 204], and cerebral blood perfusion [54, 242]. Similarly, age-dependent trajectories are well-characterized for peripheral biomarkers, including body mass index (BMI), waist circumference, blood pressure, cholesterol, creatinine, albumin, bilirubin, glucose, glycosylated hemoglobin (HbA1c), high- and low-density lipoproteins (HDL and LDL) and endocrine measures such as testosterone and follicle-stimulating hormone [3, 9, 33, 148, 170, 186, 235, 239]. An emerging question is how these brain- and body-level measurements relate to each other.

Multi-modal neuroimaging is increasingly used to quantify the effect of typical aging and neurological disease on multiple tissue types and biological systems [162]. For instance, blood perfusion reduction and vascular dysregulation are linked with greater dementia risk, and can precede observable brain atrophy and clinical symptom onset of the disease by years [72, 94, 100, 101, 233, 237]. Nevertheless, routine clinical evaluation still relies more heavily on demographic information (age and sex) and laboratory-based assessments of blood and urine panels than on neuroimaging workup. Structural magnetic resonance imaging (MRI) (T1– and T2–weighted MRI) is used in occasional relevant scenarios—to rule out possibility of focal neurological lesion such as primary or metastatic tumor or to indicate the presence of dominant atrophy—while more advanced MR imaging techniques such as functional MRI (fMRI), arterial spin labeling (ASL), and diffusion MRI feature less prominently in clinical decision-making.

Recognizing the potential for peripheral biomarkers to serve as readily accessible indicators of brain health, there is greater effort to identify translatable brain-body relationships. Initial efforts have focused on isolating specific brain-body associations between individual features of interest. Notable examples include relationships between reduced glucose tolerance and hippocampal atrophy [37], between obesity and frontal cortex thinning and subcortical volume reduction [107, 127, 222], and between hypertension and reductions in blood perfusion [6] and cortical thinning [70]. However, most of these studies focus on single brain measurements or a compact set of physiological markers. Without incorporating comprehensive measurements from both neural and peripheral systems, we cannot fully compare the sensitivity of brain features to particular physiological signatures.

Here we ask two questions. First, what can be inferred about brain health from individuals’ demographic and laboratory profiles? Second, among various MRI-derived brain measurements— spanning structural, functional, and vascular domains—which are most closely associated with specific physiological phenotypes? We address these questions using partial least squares (PLS) multivariate pattern learning to assess the covariance between multi-modal MRI brain measurements and a broad set of demographics and physiological characteristics [110, 132, 133]. We analyze two datasets with comprehensive neuroimaging and peripheral multi-omics: the Human Connectome Project–Aging [22, 82, 194] and UK Biobank [138]. We identify two generalizable axes of brain–body associations in both datasets and in both biological sexes (males and females): an aging axis and a metabolic axis. Furthermore, we identify brain features that contribute most to each axis. Finally, we examine how an individual’s position along these axes relates to their cognitive performance.

## RESULTS

The discovery dataset is obtained from the Human Connectome Project–Aging (HCP–A; https://www.humanconnectome.org/study/hcp-lifespan-aging). HCP–A data comes from 597 participants (329 females; 268 males) aged 36 to 100 years [22, 82, 194]. Detailed information on participants’ demographics, data acquisition, and preprocessing is provided in *Methods*.

We use partial least squares (PLS) [110, 132, 133] to assess how participants’ age, blood pressure, body mass index (BMI) and plasma measurements (biomarkers; Fig. S1 and Table. S1) covary with brain measurements derived from T1/T2–weighted structural imaging, arterial spin labeling (ASL) imaging, diffusion and functional magnetic resonance imaging. This multivariate analysis identified two statistically significant (assessed by permutation testing) and cross-validated latent variables (LVs) linking biomarkers to brain measurements in both males and females. In the following sections, we discuss these latent variables in detail. Throughout this report, we stratified the analysis by biological sex. There exist known differences between males and females in both brain and biomarker measures. For example, females typically exhibit higher cerebral blood perfusion than males [1, 4, 46, 53, 54, 77, 78, 102, 175, 189], along with distinct hormonal trajectories and notable differences in certain plasma markers (e.g., creatinine, and lipid markers) [13, 38, 87, 95, 112]. Therefore, we stratified the analysis by biological sex throughout the report (see Fig. S1).

Finally, we use the UK Biobank dataset (https://www.ukbiobank.ac.uk) as the validation dataset to replicate the findings. UK Biobank data comprises 3 013 participants (1 582 females; 1 431 males) aged 51 to 83 years [138, 157]. The biomarkers from this cohort are shown in Fig. S2 and Table. S2.

### The aging axis

The first latent variable (LV–I) captures 66.74% and 71.90% of the covariance between biomarkers and brain measurements in males and females, respectively (*p <* 9.99 × 10^−3^ for both groups). LV–I is characterized by a biomarker profile composed of: lower age (the strongest contributor to LV–I), alongside lower levels of follicle-stimulating hormone (FSH), urea, luteinizing hormone (LH), vitamin D, creatinine, glycosylated hemoglobin (HbA1c), and mean arterial pressure (MAP) (Fig. 1a). This biomarker profile is prominently associated with greater cortical thickness and faster cortical arterial transit time (Fig. 1b and Fig. S3). Additional associations include greater fractional anisotropy (FA), cortical functional connectivity (FC) strength, and blood perfusion, and lower mean diffusivity (MD) and white matter hyperintensity (WMH). Cortical myelin content (quantified as T1/T2 ratio) and structural connectivity (SC) strength show relatively smaller contributions (Fig. 1b and Fig. S3).

**Figure 1.**
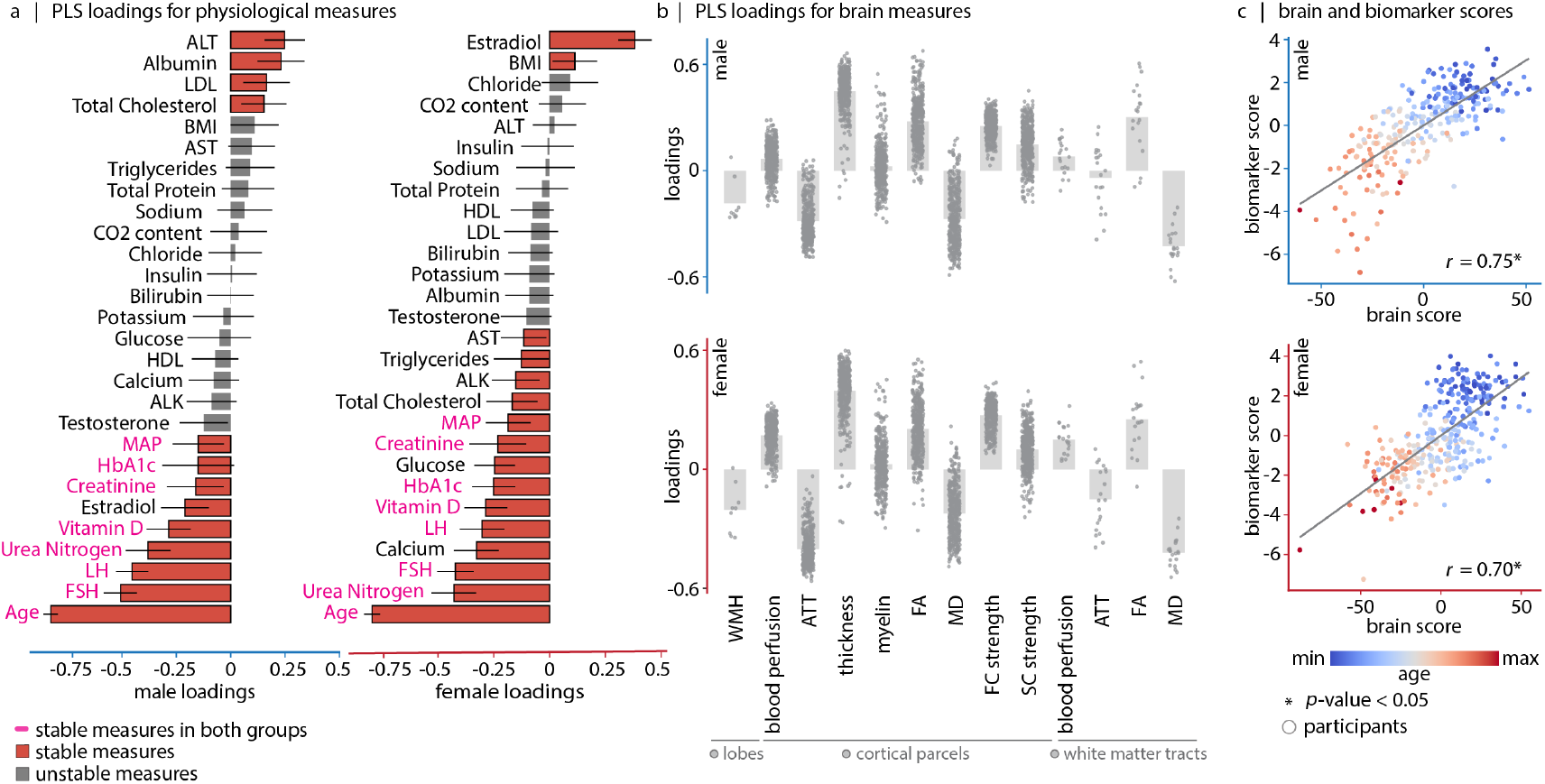
Mapping biomarkers to brain features: first latent variable (LV–I) captures the aging axis. Using PLS analysis, we identify a significant latent variable that accounts for 66.74% (males) and 71.90% (females) of the covariance between brain measurements and biomarkers—including demographics, BMI, blood pressure and blood panel measurements. (a) Biomarker loadings. Bootstrap resampling is used to estimate the stability of each individual biomarker’s contribution to the overall multivariate pattern. Each biomarker loading is divided by its bootstrap-estimated standard error, yielding a measure called “bootstrap ratio”. Bootstrap ratios are high for biomarkers with large weights (i.e., large contributions) and small standard errors (i.e., highly stable). Stable biomarkers for which the estimated 95% confidence interval do not cross zero, are shown in red. (b) Brain loadings. Each dot represents a cortical brain region as defined by the Schaefer-400 parcellation [182] or a white matter tract defined by the JHU atlas [89, 226]. For WMH, gray dots represent quantified WMH in cortical lobes and the total WMH burden. (c) Correlation between brain (*x*-axis) and biomarker scores (*y*-axis) for males (top; *r* = 0.75) and females (bottom; *r* = 0.70). Each dot represents an individual participant, colored by their age. The score per participant shows the extent to which the participant expresses the brain-biomarker association captured by LV–I. Score correlation values passed cross-validation in both sex groups.

Given that age is the strongest contributor to the LV–I pattern, we interpret LV–I as primarily capturing the “aging axis”. Other stable biomarker contributors align with age-related physiological changes (see Fig. S1). Blood urea and creatinine levels increase with age due to reduced glomerular filtration rate [122, 149, 223]. FSH and LH increase with age in males [8] and in females as a result of age-related alterations in gonadal function [114] and in the hypothalamic-pituitary unit. HbA1c level increases with age independently of elevation in glucose level and insulin resistance [50]—partly as a result of age-related reduction in red blood cell (RBC) turnover and increased RBC lifetime; the prolonged RBC lifespan extends hemoglobin exposure to circulating glucose and enhances non-enzymatic glycation of hemoglobin, creating an age-dependent elevation in HbA1c [238]. MAP increases with age due to changes in peripheral vascular resistance and artery stiffness [61]. Fig. 1c shows participants’ biomarker versus brain PLS scores, color-coded by participants’ age. In the PLS context, biomarker and brain scores quantify how strongly each participant expresses the brain-biomarker association captured by the latent variable loading profiles. Participants’ brain and biomarker scores along LV–I dimension are correlated (*r* = 0.75 for males and *r* = 0.70 for females; *p <* 9.99 × 10^−4^ for both). The correspondence between biomarker and brain scores is crossvalidated and is generalizable across participants in each biological sex group (*p <* 9.99 × 10^−3^ for both groups).

Fig. 2 shows the loading patterns for each brain measurement, and the pair-wise spatial similarity between these patterns along with their statistical significance after accounting for spatial autocorrelation. In both biological sex groups, cortical thickness and cortical mean diffusivity (MD) loadings are negatively correlated; cortical MD and fractional anisotropy (FA) loadings are also negatively correlated. Together, these findings recapitulate the previously reported brain structural deterioration with age—age ↑, cortical thickness ↓, MD ↑, FA ↓. Notably, MD and ATT loadings are aligned; indicating that with age, brain regions with prolonged arterial transit time also have greater water diffusivity— age ↑, MD ↑, ATT ↑. This coordinated pattern suggests that microstructural and vascular alterations in aging are interrelated.

**Figure 2.**
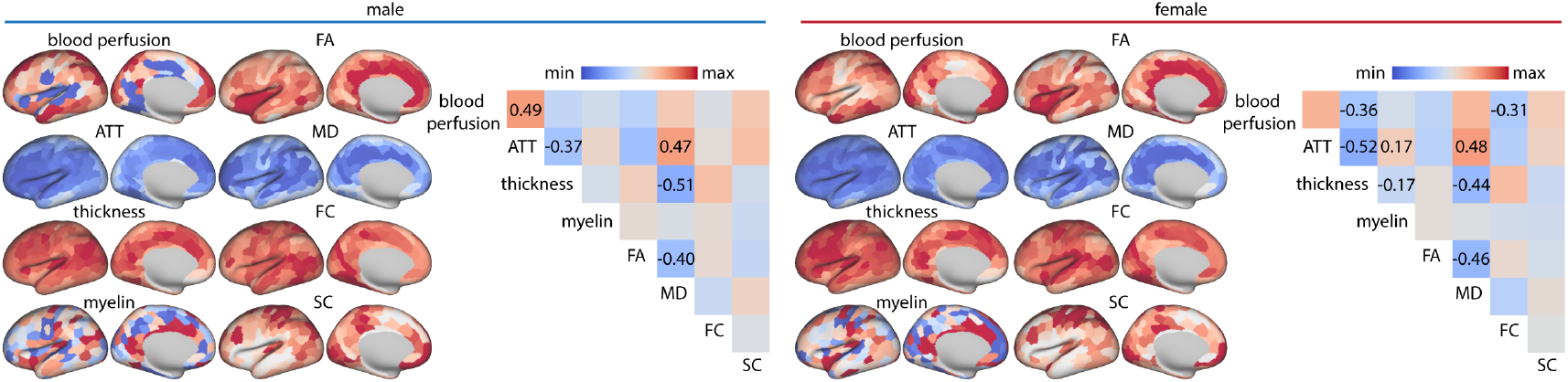
LV–I brain loadings. Left hemisphere brain loadings for each included brain feature. Participants who more strongly express the biomarker pattern shown in Fig. 1a (participants with lower age) have higher cortical thickness and blood perfusion, and lower ATT. Brain maps are shown on the fsLR inflated cortical surfaces. Bilateral hemisphere data are shown in Fig. S4. The heatmaps show similarity of brain loadings across included brain measurements. Associations that remain significant after controlling for spatial autocorrelation and multiple comparison correction—using false discovery rate (FDR)—are marked with their corresponding correlation values (ATT and MD: *p*_spin_ = 1.68 × 10^−2^ (males), *p*_spin_ = 7.99 × 10^−3^ (females); FA and MD: *p*_spin_ = 1.68 × 10^−2^ (males), *p*_spin_ = 6.99 × 10^−3^ (females); MD and thickness: *p*_spin_ = 1.68 × 10^−2^ (males), *p*_spin_ = 6.99 × 10^−3^ (females); ATT and thickness: *p*_spin_ = 1.68 × 10^−2^ (males), *p*_spin_ = 6.99 × 10^−3^ (females); perfusion and ATT: *p*_spin_ = 1.68 × 10^−2^ (males); perfusion and thickness: *p*_spin_ = 7.99 × 10^−3^ (females); perfusion and FC: *p*_spin_ = 0.014 (females); ATT and myelin: *p*_spin_ = 7.99 × 10^−3^ (females); thickness and myelin: *p*_spin_ = 6.99 × 10^−3^ (females)). In Fig. S5, we compare the aging-axis brain loading patterns between males and females. Across all included metrics, the correlation between brain loadings in males and females exceeds 0.72, indicating strong concordance in age-related brain changes between sexes.

### The metabolic axis

The second latent variable (LV–II) captures 12.83% and 14.14% of the covariance between biomarkers and brain measurements in males and females, respectively (*p <* 9.99 × 10^−4^ for both). LV–II is characterized by a biomarker profile composed of: lower BMI, MAP, HbA1c, glucose, insulin, alanine aminotransferase (ALT), as well as higher high-density lipoprotein (HDL) (Fig. 3a). On the brain side, this biomarker pattern is prominently associated with higher blood perfusion. Additional associations include greater FC strength and MD; the inverse association between BMI and MD potentially reflects the presence of gliosis [215]. Other cortical features make relatively smaller contributions (Fig. 3b and Fig. S6).

**Figure 3.**
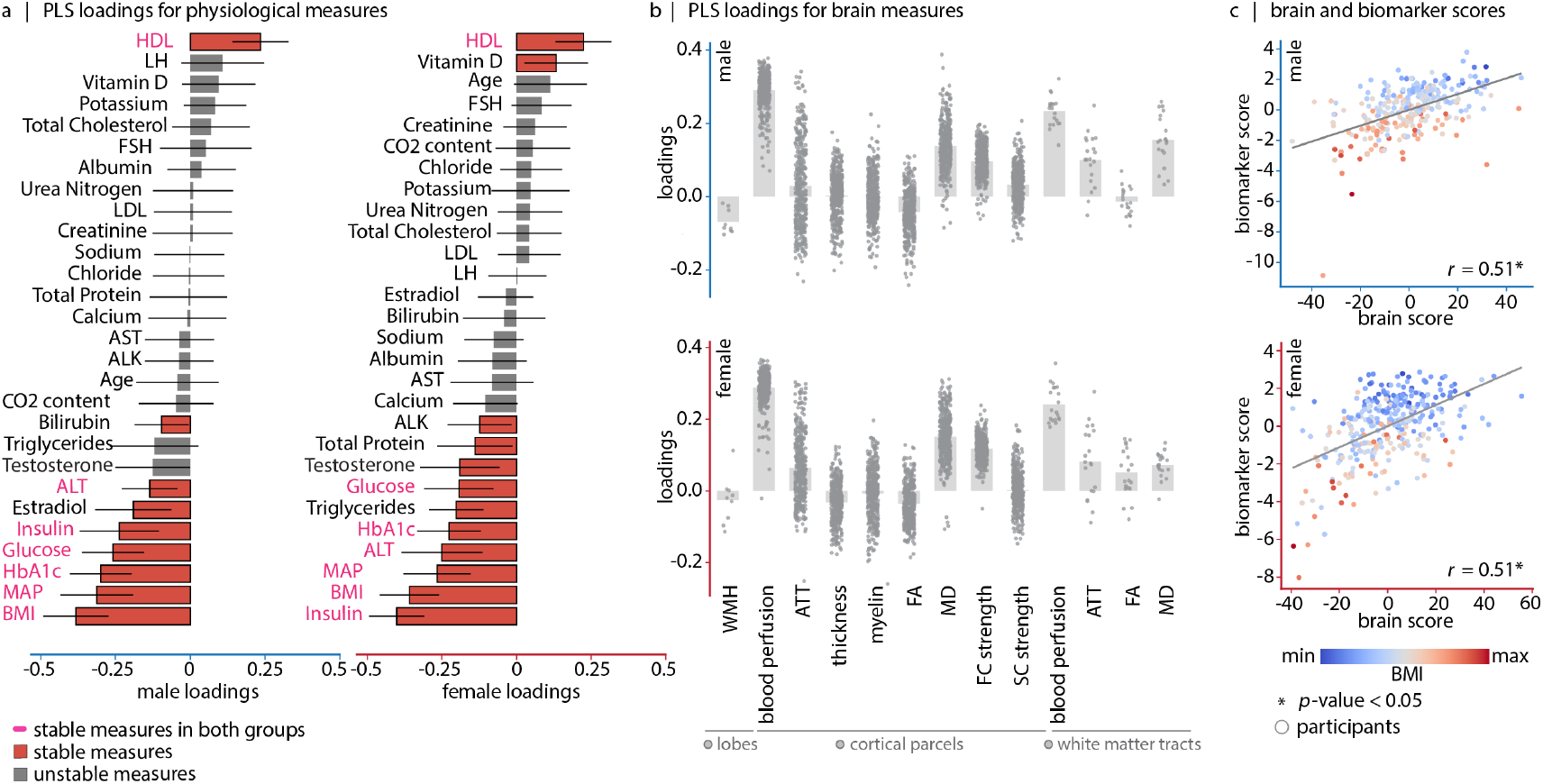
Mapping biomarkers to brain features: second latent variable (LV–II) captures the metabolic axis. Using PLS analysis, we identify a second significant latent variable that accounts for 12.83% (males) and 14.14% (females) of the covariance between brain measurements and biomarkers—including demographics, BMI, blood pressure and blood panel measurements. (a) Biomarker loadings. Bootstrap resampling is used to estimate the stability of each individual biomarker’s contribution to the overall multivariate pattern. Each biomarker loading is divided by its bootstrap-estimated standard error, yielding a measure called “bootstrap ratio”. Bootstrap ratios are high for biomarkers with large weights (i.e., large contributions) and small standard errors (i.e., highly stable). Stable biomarkers for which the estimated 95% confidence interval do not cross zero, are shown in red. (b) Brain loadings. Each dot represents a cortical brain region as defined by the Schaefer-400 parcellation [182], or a white matter tract defined by the JHU atlas [89, 226]. For WMH, gray dots represent quantified WMH in cortical lobes and the total WMH burden. (c) Correlation between brain (*x*-axis) and biomarker scores (*y*-axis) for males and females (in both cases: *r* = 0.51). Each dot represents an individual participant, colored by their BMI. The score per participant shows the extent to which the participant expresses the brain-biomarker association captured by LV–II. Score correlation values passed cross-validation in both sex groups. both males and females). The correspondence between biomarker and brain scores is cross-validated and is generalizable in each biological sex group (*p <* 9.99 *×* 10^−3^ for both groups).

The LV–II biomarker pattern can be broadly interpreted as the “metabolic axis”. Biomarker contributors to this latent variable are highly interconnected. Excessive weight (high BMI) increases activity of sympathetic nervous system and renin–angiotensin–aldosterone system, leading to elevated blood pressure [163, 196, 207]. High BMI is also linked to insulin resistance—a biological phenomenon where cells become less responsive to insulin for glucose transport [103]. Insulin resistance can be indexed by the homeostatic model assessment of insulin resistance (HOMA-IR), derived from fasting insulin and glucose concentration levels (their product divided by a constant) [128]. At the molecular level, insulin resistance develops through the accumulation of intracellular lipid metabolites, namely diacylglycerol and ceramide, which activate lipid-dependent serine/threonine kinase (including protein kinase C) and impair insulin signaling pathways—through phosphorylation of insulin receptor substrates. Insulin resistance causes impaired insulin-mediated inhibition of lipolysis, leading to increased adipose tissue breakdown and elevated circulating free fatty acids (FFAs) that further inhibit the antilipolytic effect of insulin. Lipotoxic stress and elevated FFAs affect liver function and result in nonalcoholic fatty liver disease, elevated ALT, and systemic inflammation [124, 146]. Hyperinsulinemia leads to chronic hyperglycemia and increases type 2 diabetes mellitus risk through progressive pancreatic *β*-cell exhaustion [165]. Moreover, insulin resistance contributes to endothelial dysfunction and hypertension by impairing the PI3K/Akt pathway responsible for nitric oxide-mediated vasodilation while preserving the MAPK pathway that promotes vasoconstriction through endothelin-1 production. Elevated FFAs further contribute to vasoconstriction through direct effects on vascular smooth muscle cells [147, 188, 199]. Hyperinsulinemia downregulates muscle lipoprotein lipase, reducing HDL generation from very-low-density lipoprotein processing and impairs insulin’s normal suppression of hepatic lipase, leading to increased breakdown of existing HDL particles and their rapid clearance from circulation [105]. Meanwhile, BMI has an inverse relationship with HDL—this relationship does not extend to low-density lipoprotein (LDL) and total cholesterol levels [171]. Fig. 3c shows participants’ biomarker versus brain PLS scores, color-coded by participants’ BMI. Participants’ brain and biomarker scores along this latent dimension are correlated (*r* = 0.51 for both males and females). The correspondence between biomarker and brain scores is cross-validated and is generalizable in each biological sex group (*p <* 9.99 *×* 10^−3^ for both groups).

Fig. 4 shows the loading patterns for each brain measurement, and the pair-wise spatial similarity between these patterns along with their statistical significance after accounting for spatial autocorrelation. In both biological sex groups, the blood perfusion and FC strength loadings are positively aligned, suggesting that metabolic-related reduction in perfusion is also reflected as reductions in FC strength of the same regions—metabolic dysfunction ↑, blood perfusion ↓, FC strength ↓. The third PLS latent variable did not pass cross-validation in either biological sex group (*p* = 0.16, 0.21 for males and females, respectively) and is therefore not described.

**Figure 4.**
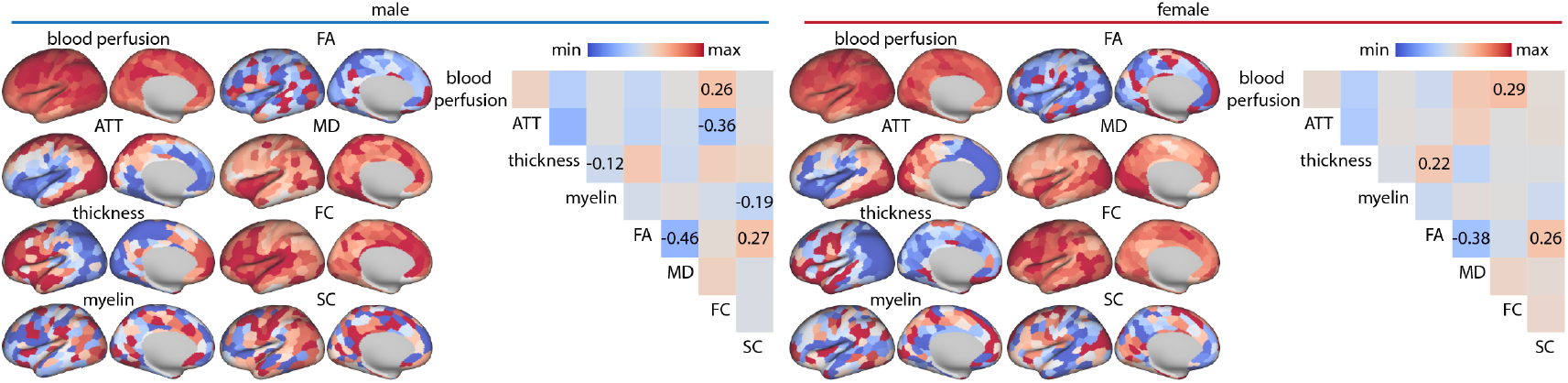
LV–II brain loadings. Left hemisphere brain loadings for each included brain feature. Participants who more strongly express the biomarkers’ pattern illustrated in Fig. 3a (e.g. those who lower BMI) have higher blood perfusion. Brain maps are shown on the fsLR inflated cortical surfaces. Bilateral hemisphere data are shown in Fig. S7. The heatmaps show similarity of brain loadings across included brain measurements. Associations that remain significant after controlling for spatial autocorrelation and multiple comparison correction—using false discovery rate (FDR)—are marked with their corresponding correlation values (FC and blood perfusion: *p*_spin_ = 2.24 × 10^−2^ (males), *p*_spin_ = 0.014 (females); FA and MD: *p*_spin_ = 6.99 × 10^−3^ (males), *p*_spin_ = 0.014 (females); FA and SC: *p*_spin_ = 6.99 × 10^−3^ (males), *p*_spin_ = 0.014 (females); myelin and SC: *p*_spin_ = 6.99 × 10^−3^ (males); thickness and myelin: *p*_spin_ = 4.66 × 10^−2^ (males); ATT and FC: *p*_spin_ = 6.99 × 10^−3^ (males); thickness and FA: *p*_spin_ = 0.014 (females)). In Fig. S8, we compare the metabolic-axis brain loading patterns between males and females.

### The metabolic axis is independent of age

We next assess whether the LV–II pattern—capturing the metabolic axis—is age-driven. To test this, we regress out the non-linear effect of age from all features included in the PLS model—biomarkers and brain measurements—using generalized additive models for location scale and shape (GAMLSS) [198]. We then rebuild the PLS model using age-corrected features. In this updated model, LV–I^*′*^ captures 34.91% and 46.15% of the covariance between age-corrected biomarkers and age-corrected brain measurements in males and females, respectively (*p <* 9.99 × 10^−4^ for both). The LV–I^*′*^ pattern is shown in Fig. 5 and it resembles LV–II from the original PLS model (based on uncorrected data; Fig. 3), albeit with some differences (e.g., in males, vitamin D becomes a stable contributor; in females, vitamin D is no longer stable, while calcium becomes stable). This latent variable is cross-validated in both sex groups (*p <* 9.99 *×* 10^−3^ for both groups).

**Figure 5.**
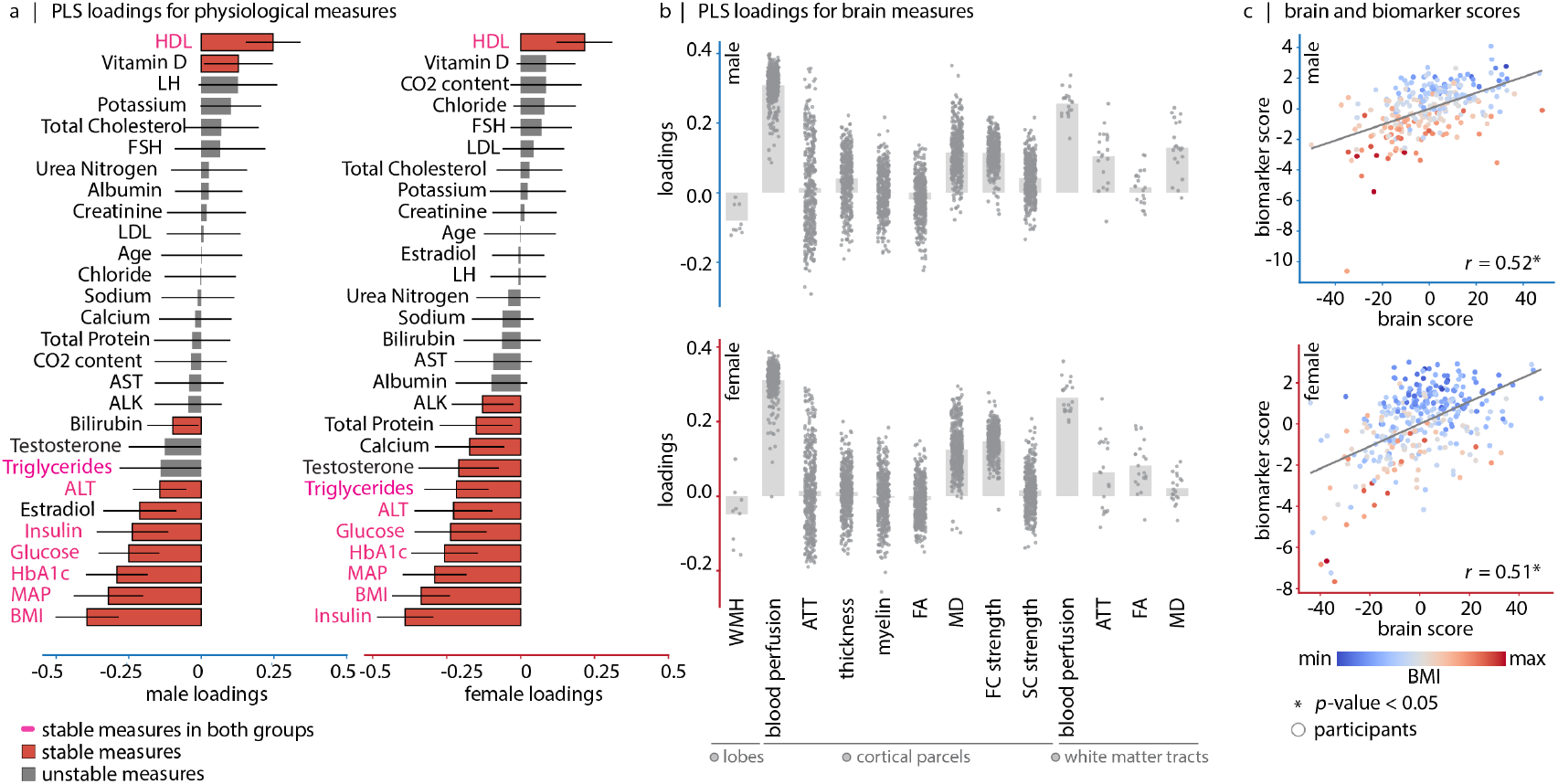
Mapping biomarkers to brain features after regressing out the age effect: first latent variable (LV–I^*′*^) captures the metabolic axis. We use GAMLSS models to regress out the age effect from all included features prior to the implementation of the PLS model (age is regressed out from both brain and biomarker measures). We identify a significant latent variable that accounts for 34.91% and 46.15% of the covariance in the data for males and females, respectively. (a) Biomarker loadings. Bootstrap resampling is used to estimate the stability of each individual biomarker’s contribution to the overall multivariate pattern. Each biomarker loading is divided by its bootstrap-estimated standard error, yielding a measure called “bootstrap ratio”. Bootstrap ratio is high for biomarkers with large weights (i.e., large contributions) and small standard errors (i.e., highly stable). Stable biomarkers for which the estimated 95% confidence interval do not cross zero, are shown in red. (b) Brain loadings. Each dot represents a cortical brain region as defined by the Schaefer-400 parcellation [182], or a white matter tract defined by the JHU atlas [89, 226]. For WMH, gray dots represent quantified WMH in cortical lobes and the total WMH burden. See Fig. S10 for brain loadings shown on the fsLR cortical surface. (c) Correlation between brain (*x*-axis) and biomarker scores (*y*-axis) for males (top; *r* = 0.52) and females (bottom; *r* = 0.51). Each dot represents an individual participant, colored by their BMI. The score per participant shows the extent to which the participant is expressing the brain-biomarker association captured by LV–I^*′*^.

This first latent variable highlights the link between poor body metabolic profile (characterized by higher BMI, MAP, HbA1c, glucose, insulin and ALT, and lower HDL) and lower brain blood perfusion. The spatial pattern of reduction in blood perfusion aligns with the FC strength reduction pattern (Fig. S10). Given that the most extreme stable biomarker loadings are similar across males and females, we further assess the similarity of brain loadings across sexes. We find the highest correspondence for ATT (*r* = 0.82, FDR-corrected *p*_spin_ = 1.59 × 10^−3^) and blood perfusion maps (*r* = 0.63, FDR-corrected *p*_spin_ = 1.59 × 10^−3^), respectively. Structural connectome strength (*r* = 0.06, FDR-corrected *p*_spin_ *>* 0.05), FA (*r* = 0.20, FDRcorrected *p*_spin_ = 2.66 × 10^−3^), and myelin maps (*r* = 0.26, FDR-corrected *p*_spin_ = 1.59 × 10^−3^) exhibit lower cross-sex spatial correspondence (Fig. S11). In Fig. S9 we show the superiority of blood perfusion measure over other functional and structural brain imaging measurements in capturing the potential effects of worse metabolic index. Taken together, these findings indicate that the captured metabolic axis reflects a robust pattern beyond agerelated variations. Rather, it reflects a distinct link between systemic metabolic dysfunction and vascular and functional alterations in the brain.

### Cognitive relevance of aging and metabolic dysfunction

What are the behavioral manifestations of the captured brain-body interaction axes in the typical aging cohort? The HCP–A dataset provides a rich array of neuroimaging, demographic, physiological and cognitive assessments that allow us to address this question. Here, we examine the link between a range of cognitive measures (fluid, crystallized and total cognition composite scores) and participants’ brain scores on the “aging” and “metabolic” axes.

Participants—both males and females—with greater expression of brain patterns shown in Fig. 2a (brain LV–I pattern; younger age) have greater cognitive flexibility and higher total cognition scores. Meanwhile, they perform worse on crystallized cognition tasks, indicating weaker performance in abilities heavily influenced by prior learning (Fig. 6a). Female participants with greater expression of multivariate brain patterns shown in Fig. 5a—characterized by healthier metabolic status—also express cognitive differences compared to their less metabolically healthy female counterparts and perform better in tasks requiring cognitive flexibility (Fig. 6b). This finding aligns with reported sex differences in the association between metabolic syndrome and dementia, with metabolic syndrome being significantly associated with cognitive impairment in women only [34, 35, 42, 113, 129, 184]. In short, fundamental physiological and neuroimaging indicators of metabolic status manifest in individual differences in cognitive performance.

**Figure 6.**
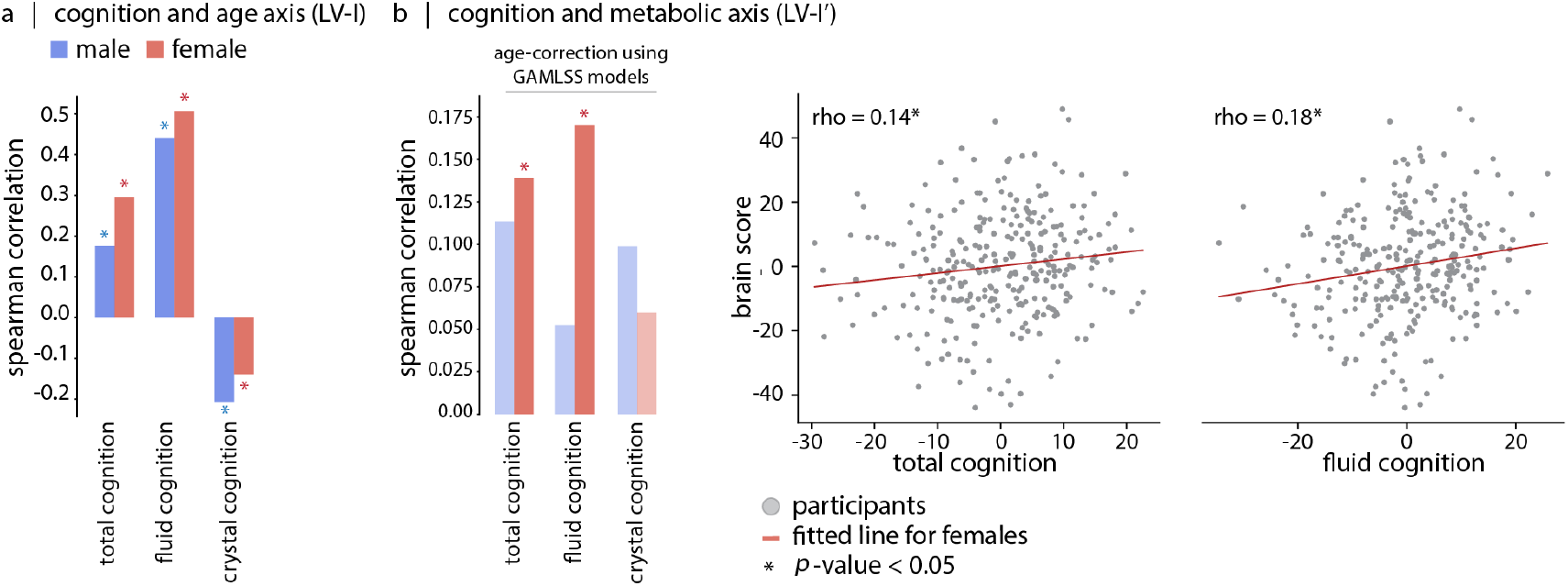
Cognitive relevance of aging and metabolic axes. (a) Spearman’s rank correlation between LV–I brain scores and total, fluid, and crystallized cognition composite scores (total cognition: *ρ* = 0.18, *p*_perm_ = 7.99 × 10^−3^ (males), *ρ* = 0.30, *p*_perm_ = 1.49 × 10^−3^ (females); fluid cognition: *ρ* = 0.44, *p*_perm_ = 2.99 × 10^−3^ (males), *ρ* = 0.51, *p*_perm_ = 1.49 × 10^−3^ (females); crystallized cognition: *ρ* = 0.21, *p*_perm_ = 4.49 × 10^−3^ (males), *ρ* = 0.14, *p*_perm_ = 1.69 × 10^−2^ (females)). (b) Spearman’s rank correlation between LV–I^*′*^ brain score and cognitive measures, while GAMLSS models are used to correct for non-linear age effects in the feature set. In this case, LV–I^*′*^ is capturing the metabolic axis (see Fig. 5). Fluid and total cognition composite scores are associated with brain scores in females (total cognition: *ρ* = 0.14, *p*_perm_ = 2.55 × 10^−2^ (females); fluid cognition: *ρ* = 0.18, *p*_perm_ = 8.99 × 10^−3^ (females)). Reported *p*-values are corrected for multiple comparison using FDR within each biological sex-group.

### Validation: aging and metabolic axes in UK Biobank

To validate the findings, we use the UK Biobank dataset (https://www.ukbiobank.ac.uk; see *Methods*). We use PLS to assess how plasma and urine panel measures, age, blood pressure, BMI, and various anthropometric measurements (hip, waist, and body fat percentage) covary with a series of brain measures (see Fig. S2) [52]. The assessed brain measurements include ASL-derived ATT and blood perfusion, cortical thickness, area and volume of brain regions in addition to FA, MD, intra-cellular volume fraction (ICVF), isotropic volume fraction (ISOVF) of white matter tracts and quantified whole brain white matter hyperintensity (WMH) volume. Imaging acquisition parameters, parcellations (Desikan-Killiany atlas for cortex) and preprocessing pipelines for the UK Biobank dataset are all different from those used in the HCP– A dataset. In the following paragraphs, we discuss the two first statistically significant and crossvalidated LVs for both males and females.

The first latent variable captures 75.71% and 78.11% of the covariance between biomarkers and brain measurements in males and females, respectively (*p*_perm_ *<* 9.99 × 10^−4^ for both). The biomarker profile for LV–I is prominently driven by age (Fig. S12a), indicating that aging is associated with higher ATT and WMH volume, lower blood perfusion, lower cortical and subcortical volumes, lower cortical areas and thickness, in addition to lower white matter FA, higher white matter MD, lower ICVF, and higher ISOVF (Fig. S12b and Fig. S13). This pattern is similar to the “aging axis” presented at Fig. 1. The directionality of microstructural changes found here are aligned with age-associations in the UK Biobank dataset previously described by Cox et al. [40].

The second latent variable captures 15.33% and 14.93% of the covariance between biomarkers and brain measurements in males and females, respectively (*p <* 9.99 × 10^−4^ for both). The biomarker profile for LV–II captures the metabolic dysfunction profile (Fig. S14a). LV–II biomarker profile is highlighting low HDL and age, along with high BMI, hip and waist circumferences, body fat percentage, diastolic blood pressure, ALT, aspartate aminotransferase (AST), triglyceride, c-reactive protein—previously linked to metabolic syndrome and inflammatory disorders [227]—and sodium in urine. This metabolic profile is dominantly associated with lower brain blood perfusion (Fig. S14b and Fig. S15).

We further show that the LV–II (metabolic axis) from UK Biobank is not age-driven. We use GAMLSS models to regress out the non-linear age effects (as well as the site effect) from all biomarkers and brain features; we then rebuild the PLS model. Fig. S16a,b shows the first latent variable, which again resembles the biomarker profile associated with metabolic dysfunction and highlights its relevance to blood perfusion (also see Fig. S17). Last, we examine how brain loadings from LV–I^*′*^ (after correcting for age) relate to participants’ fluid intelligence. In both female and male groups from the UK Biobank dataset, metabolically healthier individuals demonstrated better fluid intelligence scores (*ρ* = 0.07, *p*_perm_ = 1.69 *×* 10^−2^ (males), *ρ* = 0.10, *p*_perm_ = 9.99 *×* 10^−3^ (females)) (Fig. S18).

## DISCUSSION

In the present report, we analyze data of HCP–Aging and UK Biobank cohorts to characterize the multivariate relationships between a series of brain measurements and bodily physiological phenomics— including age, body mass index (BMI), blood pressure, and plasma/urine protein profiles [22, 82, 138, 157, 194]. The PLS analysis [110, 132, 133] reveals two principal axes of brain-body associations: one driven primarily by age, and the other by metabolic health. The most prominent axis of association is age-related. In both datasets, increased age is associated with cortical thinning and microstructural alteration, along with slower arterial transit time and reduced cerebral blood perfusion. The second axis of brain-body covariance is related to metabolic features. Elevated BMI, blood pressure, HbA1c, glucose, and ALT liver enzyme, along with lower HDL cholesterol are linked to lower brain blood perfusion. We further show that the captured metabolic effect on the brain is accompanied by lower cognitive flexibility; reflecting the potential behavioral consequences of the observed brain-body relationships.

Aging is a time-dependent biological phenomenon that affects every system in the human body [121], from the skeletal and muscular to the cardiovascular and nervous systems. Previous research on brain aging has established typical aging trajectories for various neuroimaging measures that serve as benchmarks to detect pathological deviations from the norm [20]. Brain age-related changes are now documented across multiple brain features, including cortical thickness [48, 59, 60, 86, 131, 195, 206], gyrification [86], brain tissue volume [20, 193, 232], FA [18, 73, 88, 109, 202], MD [18, 28, 88, 96, 232], SC [45, 123, 245], FC [19, 30, 45, 187, 197, 204, 211], and blood perfusion [54, 102, 242]. Across these modalities, there is growing consensus regarding age-related structural deterioration and reorganization of brain vascular and functional networks.

PLS brain loadings for the aging axis show consistent directionality in both datasets, reflecting brain structural deterioration alongside cerebrovascular decline. Specifically, we recapitulate agerelated structural deterioration, characterized by cortical thickness ↓, FA ↓, MD ↑, ICVF ↓, ISOVF ↑, as well as cerebrovascular dysfunction, characterized by blood perfusion ↓, FC strength ↓, WMH ↑, ATT ↑. However, while cortical thickness shows the greatest loading magnitude (e.g., most affected by aging) in HCP–Aging, blood perfusion, transit time, and white matter hyperintensity loadings dominate in UK Biobank. The differences in brain loading ranking may originate from distinct age ranges in HCP– Aging (36–100 years) and UK Biobank (51–83 years) cohorts, or from their imaging protocol differences.

Our study resonates with a growing appreciation for studying brain aging from a wider, multivariate perspective, and is in line with the studies which suggest superior brain-age prediction accuracy when incorporating multi-modal brain imaging— T1–weighted MRI, T2–FLAIR, T2*, diffusion-MRI, task and resting-state fMRI—compared to single-modality approaches [36, 99, 116, 152, 177]. Our findings emphasize that the spatial patterns of agerelated changes are not identical across “all” brain features. For example, while FA and MD have spatially similar age-related profiles, blood perfusion exhibited a distinct pattern of age-related decline; suggesting that different neuroimaging measures capture complementary aspects of brain aging. We further show that age-related brain loadings are relatively consistent across biological sex groups.

One consistent and reproducible finding across this study is the identification of a metabolic axis linking peripheral physiological markers and the brain. Prior studies have shown that metabolic factors are associated with brain health; for example, increased BMI and abdominal fat have been linked to both functional [161] and structural brain alterations [25, 39, 47, 62, 63, 106, 111, 127, 160, 169, 217, 222, 234]; and increased HbA1c and adiposity-related insulin resistance is linked to cortical thickness changes [190, 191]. Extending beyond these previously documented associations, we show that cerebral blood perfusion is the primary brain feature that potentially declines as a function of elevated metabolic risk factors. One possibility is that vascular susceptibility to metabolic risks constitutes the principal biological mechanism underlying the brain-body metabolic axis in aging populations. Elevated LDL cholesterol [26, 57, 69], glucose, glycosylated hemoglobin [23], blood pressure [150, 183, 192], body fat and possibly BMI [130] as well as low HDL cholesterol [12] increase atherosclerotic and cardiovascular disease risk, compromise vascular integrity, and ultimately impair perfusion.

Various biological pathways have been suggested to explain how metabolic abnormalities may disrupt endothelial function. For instance, HDL is known to have anti-inflammatory and antioxidant properties [225], by mediating cholesterol trafficking and reducing oxidized lipids from LDL, inhibiting endothelial apoptosis and stimulating their migration, and preventing monocytes adhesion to the endothelial surface—a critical step in the development of atherosclerotic plaques [15, 139, 213]. When HDL levels are low, vascular resilience is reduced, leaving the brain particularly vulnerable to vascular injury. Indeed, low serum HDL is linked to compromised blood-brain barrier integrity, increased cerebral amyloid burden [173] and enhanced *β*-amyloid accumulation within cerebral vessels [29]; there are also studies that report association of lower HDL (and ApoAI—the major protein component of HDL) with Alzheimer’s disease and individual cognitive performance [17, 29, 117, 136, 174, 247]. Taken together, a healthier metabolism may foster vascular resilience and vascular reactivity and sustain healthy cerebral blood perfusion, which further protects against neurodegeneration.

The perfusion-centric metabolic effect can partly explain the disparate structural effects of metabolic risk factors reported to date [135]. While most studies report reductions in frontal cortex volume and thickness in obesity [25, 39, 62, 63, 111, 127, 160, 234], some report overall gray matter loss or even white and gray matter expansion [80, 107, 217]. In our PLS model, the metabolic axis is character-10 ized by lower frontal cortex thickness and greater occipital cortex thickness in the HCP–A dataset. This pattern is consistent with independent findings [85, 143, 160, 217] and is in line with the findings of Peterson and colleagues [164], who reported a similar cortical thickness pattern associated with metabolic syndrome risk factors when analyzing data of over 40 000 UK Biobank participants. Several mechanisms are proposed for obesity- and metabolic syndrome-related cortical thickening in occipital cortex, ranging from neuroinflammation [143] to enhancement of visual processing and attention networks in obesity [217]. Given that metabolic markers covary most strongly with cerebral blood perfusion in both datasets, future neuroimaging studies of obesity and metabolic syndrome may achieve greater consistency by including blood perfusion measures when assessing brain health.

Our results also suggest a pattern of blood plasma markers and body vitals that are closely associated with brain physiology. Defining “metabolic dysfunction” remains an important challenge in the field, with ongoing debate over which features should be included in clinical definitions [75, 104, 158, 172]. Current discussions often focus on including or excluding features from the definition, often without a specific weighting of the relative importance of these plasma and urine factors in the neurologic context. For instance, one definition of metabolic dysfunction is presence of three or more of the listed elements including elevated glucose, blood pressure, triglycerides, waist circumference, and reduced HDL [180]. Our cross-dataset and cross-sex consensus suggests that higher HDL cholesterol may be neuroprotective, while elevated BMI, HbA1c, blood pressure, and ALT consistently emerge as factors associated with lower brain blood perfusion.

This study provides a framework for studying the mechanisms of neuroprotection and vulnerability in metabolic dysfunction. Some individuals demonstrate resilience to expected metabolic brain correlates despite peripheral risk factors, while some show heightened vulnerability (discrepancy between brain and peripheral biomarker profiles; see Fig. 3c, 5c, S14c and S16c). These individual differences provide an opportunity for targeted genetic investigations to identify genetic variants that may confer resilience to brain alterations caused by metabolic risks [216].

The brain-body metabolic axis may in part reflect background genetic predisposition rather than a direct causal effect of peripheral metabolic dysfunction on the brain. Most obesity-associated genes are expressed in the central nervous system, with obesity-associated loci overlapping with genes and pathways implicated in neurodevelopment [118, 240]. These genetic influences extend beyond energy homeostasis- and reward-related brain areas, and affect broadly distributed neural circuits across the brain [210]. In other words, rather than the presence of a pure “causal” relationship between metabolic dysfunction and brain features, it is possible that individuals with higher genetic risk for obesity also have genetic propensity for the brain patterns identified in this study [217]. In line with this notion, Morys and colleagues have shown that polygenic risk scores for obesity are negatively correlated with frontal cortex thickness and volume in adolescents (ABCD cohort; mean age = 10 years) [144]. This brain pattern is related to increased impulsivity and subsequent BMI increase at one-year follow-up [145]. Hence, genetic predisposition may initiate chronic obesity and cardiovascular consequences that further impact the brain in older individuals. In either case, reversible brain alterations (both functional and structural) following weight loss [137, 214, 218, 228, 241, 244] highlight the contribution of modifiable lifestyle factors in maintaining brain health. Indeed, weight loss, following either Roux-en-Y gastric bypass surgery [5] or a simple 12-months behavioral change (diet and exercise) have been shown to increase cerebral blood flow [200].

Lifestyle changes that ameliorate (or avoid) metabolic dysfunction can have a significant clinical relevance due to their influence on brain blood perfusion health. Hypoperfusion and vascular dysfunction impair the brain’s clearance of solutes and metabolites through compromised interstitial fluid drainage leading to accumulation of toxic proteins [10, 51]. Simultaneously, hypoperfusion induces oxidative stress, promoting the formation of misfolded proteins and amyloid-*β* (*Aβ*) precursors, and further reducing the expression and activity of neprilysin, a major *Aβ*-degrading enzyme [246]. These pathological processes create a vicious cycle, as *Aβ* peptides themselves can further impair vasoregulation [153] and reduce blood flow [154], perpetuating the hypoperfusion that initially contributed to their formation. Meanwhile, animal studies have showed that induced hypoperfusion can accelerate cerebral amyloid angiopathy [156]. In short, a potential mechanistic explanation is that obesity leads to cerebrovascular alterations, which in turn facilitates amyloid-*β* deposition in brain tissue (leading to Alzheimer’s disease) and amyloid formation in brain vasculature system (leading to cerebral amyloid angiopathy). Furthermore, reduced blood flow due to arterial narrowing promotes thrombosis and increases risk of emboli formation [31]. Maintaining metabolic health may thus prevent or slow down neurodegenerative cascades by preserving brain perfusion.

The present study is part of a wider effort to understand links between the brain and the body. These bidirectional interactions are increasingly explored in the context of the gut-brain axis [142] and the proteomics-brain axis [93, 155]. The gutbrain axis consists of two communication mechanisms: one mediated based on chemicals including, neurotransmitters, neuropeptides, hormones, and gut microbiome [159], and the other mediated by the physical connections via the vagus nerve. This axis is increasingly recognized as important for understanding the spread of proteinopathies in neurodegenerative diseases [181, 229]. Furthermore, there is evidence that this axis of interaction plays a role in obesity and metabolic health via serotonergic signaling [92, 98]. Concurrently, proteomics can also help to monitor brain health, and specifically to define protein signatures associated with obesity, type 2 diabetes, and neurodegenerative diseases. The identified protein profiles enable disease vulnerability predictions and identification of potential target molecules for effective intervention plans [93, 119, 155, 176, 203, 236].

Importantly, both aging and metabolic brain signatures have behavioral relevance across participants. We replicate well-known associations between aging and cognitive domains; in particular, we find the relationship between aging and diminished cognitive flexibility [41, 56, 65, 84]. In addition, participants with greater scores on the metabolic dysfunction axis show greater cognitive impairment compared to their metabolically healthier counterparts. This finding aligns with reports documenting the interplay between diabetes mellitus type 2, low cerebral blood flow and worse memory, executive function and processing speed [14], as well as studies linking cortical thickness or brain structural volume signatures of BMI and metabolic status to participants’ working memory performance [167], fluid intelligence and prospective memory [248]. Note however that there are several studies that find no significant associations between specific metabolic syndrome features and cognition [79, 212]. The present report shows that a composite metabolic brain-body profile may be a powerful way to identify links between physiological measurements and cognitive outcome.

Our results point to possible sex differences in metabolic-cognitive associations. In the HCP–A dataset, greater metabolic deviation from health is associated with lower fluid cognition scores in females, whereas no significant negative associations survives multiple comparison correction in males. Greater female cognitive susceptibility to metabolic dysregulation is also reported in multiple independent studies [34, 35, 42, 113, 129, 184]. The potentially greater cognitive effects observed in females may account for the higher Alzheimer’s disease prevalence in females compared to males [168]. Differential cognitive risk and resilience may be driven by sex-related factors ranging from physiological differences to acquired socioeconomic disparities [11, 120]. Metabolic health and sex-differences should be integrated in future aging and neurodegenerative disease studies to understand cognitive resilience disparities across individuals in greater details.

The present findings should be interpreted with respect to several methodological considerations. First, cross-sectional associations between brain measurements and peripheral biomarkers are not causal—longitudinal studies, and interventional studies—e.g., brain imaging before and after obesity-reduction trials—can help understand the nature of associations (causal or co-existence). Second, while our findings suggest that ASL imaging may offer greater sensitivity for obesity and metabolism research, the inherently low signal-to-noise ratio of this imaging modality and its sensitivity to obesity-related confounds (such as motion artifacts, different body composition, and head placement [21, 141, 205]) is an ongoing topic of research. Encouragingly, changes in FC strength and in perfusion in the presence of metabolic dysfunction are similar. Third, in the UK Biobank dataset, laboratory-based biomarkers were acquired at the baseline visit, whereas the neuroimaging data was acquired during a follow-up session (time interval between sessions can span several years, see Fig. S2). We addressed this temporal discrepancy by using GAMLSS models to regress out age effects from all features entered to the PLS analysis.

## METHODS

All code used to perform the analyses are available on GitHub at https://github.com/netneurolab/Farahani_Metabolic_Health/.

### HCP–A: Demographics

We analyzed data from 597 participants (329 females; 268 males) aged 36–100 years from the HCP– Aging dataset (HCP Lifespan studies, 2.0 Release) [22, 82, 194]. Participants represented a “typical” aging population and had common health conditions (e.g., hypertension) without identified pathological causes of cognitive decline (e.g., stroke). All study procedures were conducted in accordance with the principles expressed in the Declaration of Helsinki and were approved by the Institutional Review Board at Washington University in St. Louis.

### HCP–A: Brain imaging acquisition

All HCP–A brain imaging data was acquired using a 3.0 Tesla Prisma scanner (Siemens; Erlangen, Germany) and a 32–channel Prisma head coil [22, 82, 194].

T1–weighted structural data was acquired using a multi-echo magnetization-prepared rapid gradient echo (MPRAGE) sequence with the following parameters: repetition time (TR) = 2 500 ms, inversion time (TI) = 1 000 ms, echo times (TE) = 1.8*/*3.6*/*5.4*/*7.2 ms, spatial resolution = 0.8× 0.8 × 0.8 mm^2^, number of echoes = 4 and flip angle = 8^*°*^. T2–weighted structural data was acquired using a 3D sampling perfection with application-optimized contrasts using different flip angle evolutions (SPACE) sequence, with the same spatial resolution as the T1–weighted image. Parameters for the T2–weighted sequence were: TR = 3 200 ms, TE = 564 ms and turbo factor = 314. Both T1– and T2–weighted images captured a sagittal field of view measuring 256 × 240 × 166 mm, and a matrix size of 320 × 300 × 208 slices. Additional acquisition parameters included 7.7% slice oversampling, 2–fold in-plane acceleration (GRAPPA) in the phase encoding direction, and a 744 Hx*/*Px pixel bandwidth. T1– and T2–weighted structural data provided the anatomical reference for analysis of all imaging modalities, reconstruction of cortical surfaces, and estimation of cortical myelin content (T1/T2 ratio) and cortical thickness [82].

Diffusion data was acquired using a spin-echo echo-planar imaging (EPI) with the following parameters: TR = 3.23 s, flip angle = 52°, spatial resolution = 1.5 × 1.5 × 1.5 mm^3^, 185 directions on 2 shells, *b* = 1500*/*3000 s*/*mm^2^, along with 28 *b* = 0 s*/*mm^2^ images. This data was used to derive FA, MD, and structural connectome strength metrics.

Resting-state fMRI data with blood-oxygen-level-dependent (BOLD) contrast was acquired using a 2D multi-band gradient-recalled echo (GRE) echoplanar imaging (EPI) with the following parameters: TR/TE = 800*/*37 ms, flip angle = 52^*°*^, spatial resolution = 2.0 × 2.0 × 2.0 mm^3^. Functional scans were acquired in pairs of two runs (four runs in total per participant, each run lasting 6.5 min), with opposite phase encoding polarity so that the fMRI data in aggregate was not biased toward a particular phase encoding polarity (two runs had the phase encoding of anterior-to-posterior (AP) and two runs had the Altogether, the brain feature set included 3 289 phase encoding of posterior-to-anterior (PA)). During rs-fMRI scanning, participants viewed a small white fixation crosshair on a black background. The functional MRI minimal preprocessing steps that were applied to the data are provided by Glasser et al. [67].

Blood perfusion and arterial transit time were measured using arterial spin labeling (ASL) magnetic resonance imaging (MRI) [102, 108]. ASL data was acquired using a pseudo-continuous arterial spin labeling (pCASL) and 2D multi-band (MB) echo-planar imaging (EPI) sequence. Pseudocontinuous ASL data was acquired with labeling duration of 1 500 ms and five post-labeling delays of 200 ms, 700 ms, 1 200 ms, 1 700 ms, and 2 200 ms, containing 6, 6, 6, 10, and 15 control-label image pairs, respectively. To calibrate perfusion measurements into units of ml*/*100g*/*min, two PD–weighted M0 calibration images (TR *>* 8 s) were acquired at the end of the pCASL scan. Other sequence parameters included: spatial resolution = 2.5 × 2.5 × 2.5 mm^3^, and TR/TE = 3 580*/*18.7 ms. For susceptibility distortion correction, two phase-encoding-reversed spin-echo images were also acquired. Participants viewed a small white fixation crosshair on a black background during the scan time (5.5 min). The ASL data preprocessing was conducted following the ASL Pipeline for the Human Connectome Project available at https://github.com/physimals/HCP-asl/ (explained in detail in Kirk et al. [108]). To run this pipeline we used QuNex platform (singularity container, version 0.99.1) [97].

### HCP–A: Brain imaging measurements

To link neuroimaging measurements with peripheral biomarkers, we incorporated a comprehensive set of cortical and subcortical features. For cortex, the following measures were included: cortical thickness, myelin, fractional anisotropy (FA), mean diffusivity (MD), cerebral blood perfusion, arterial transit time (ATT), functional connectivity (FC) strength, and structural connectivity (SC) strength. All cortical measures were parcellated using the Schaefer400 atlas [182].

For white matter, we used the Johns Hopkins University (JHU, threshold 25%) atlas to extract FA, MD, blood perfusion, and arterial transit time across 20 white matter tracts [89, 226]. White matter hyperintensities (WMH) were represented by nine measures, one reflecting the total WMH voxel count and eight reflecting WMH voxel counts within each cortical lobe (frontal, temporal, parietal, and occipital), separately for the left and right hemispheres.

Altogether, the brain feature set included 3 289 values. Among the 597 participants included in the multivariate mapping between brain and biomarkers, 62 females and 63 males had missing values for WMH measurements. All missing values were imputed using the median of the corresponding sexspecific feature.

### HCP–A: Biological samples and vital signs

The HCP–A dataset included a comprehensive array of participant-specific blood biochemistry results and physiological measurements [22]. These assessments included common health indicators including total protein, glucose, insulin, glycosylated hemoglobin (HbA1c), triglycerides, lowdensity lipoprotein (LDL), high-density lipoprotein (HDL), total cholesterol, albumin, bilirubin, creatinine, urea, chloride, sodium, potassium, calcium, vitamin D, and CO_2_ content, as well as liver metabolic enzymes including alanine aminotransferase (ALT), aspartate aminotransferase (AST), and alkaline phosphatase (ALK). Multiple hormonal measures were also obtained, including serum estradiol, testosterone, luteinizing hormone (LH), and follicle-stimulating hormone (FSH). Blood samples were collected, preferably after an 8–h fasting period. Furthermore, participants’ height and weight were also recorded, allowing the calculation of body mass index (BMI). Systolic and diastolic blood pressure/pulse values were measured, from which the mean arterial pressure (MAP) was derived. Participants’ age was also provided. In total, 28 biomarkers were included in the analysis for each participant.

Among the 597 participants (329 females; 268 males) with available biochemical data, some had missing values for specific measures: 2 females for BMI, 4 females and 2 males for MAP, 3 females and one male for HbA1c, 2 males and a female for FSH, 2 males and a female for LH, 2 males and a female for Estradiol, and one female and one male for Testosterone (see Table. S1). In these instances, the missing values were imputed using the median of the respective measure (within each sex group) to prevent not-a-number values in the PLS analysis.

### HCP–A: Cognitive assessment

Cognitive performance in HCP–A participants was evaluated using the NIH Toolbox for the Assessment of Neurological and Behavioral Function (NIHTB). Crystallized, fluid, and total cognitive composite scores were obtained for all participants. Crystallized cognitive composite score captures abilities influenced by acquired knowledge and was derived from two assessments: the oral reading recognition test and the picture vocabulary test. Fluid Cognition Composite score captures processing efficiency and novel problem-solving abilities. This composite score was derived from five components, including dimensional change card sort test, list sorting working memory test, picture sequence memory test, pattern comparison processing speed test, and flanker inhibitory control and attention test. In this study, composite scores were used in place of individual test scores due to their enhanced reliability and reduced measurement error [231].

When calculating correlations between PLS scores and cognitive scores, data of individuals with missing cognitive scores were excluded. Among Females, 36 missing values existed for cognitive measures, and among males 54 missing values existed for crystallized and total cognition scores, and 53 missing values existed for fluid cognition scores.

### UK Biobank: Demographics

We analyzed data from 3 013 participants (1 582 females; 1 431 males) aged 51–83 years (at the time of brain imaging visit) from the UK Biobank dataset. We included participants who had both the required brain imaging data—specifically ASL imaging acquired at time-point 0.2, corresponding to the initial imaging session—and peripheral biomarker data. Participants were excluded if they had more than 40 missing values in the brain imaging feature set. Participants with a history of diagnosed mental health conditions were excluded. Excluded mental health conditions included depression, mania, hypomania, bipolar or manic-depression, autism spectrum disorders, panic attacks, obsessive-compulsive disorder (OCD), schizophrenia, personality disorders, and attention deficit disorders (ADD/ADHD). Participants who self-reported neurological or psychiatric illnesses were also excluded. These included Parkinson’s disease, dementia or Alzheimer’s disease, cognitive impairment, multiple sclerosis, other demyelinating diseases, stroke, transient ischemic attack (TIA), ischaemic stroke, epilepsy, brain hemorrhage, subdural or subarachnoid hemorrhage, brain abscess or intracranial abscess, encephalitis, cerebral palsy, cerebral aneurysm, meningitis, meningioma or other benign meningeal tumors, motor neuron disease, nervous system infections, neurological trauma or injury, head injury, acute infective polyneuritis (Guillain-Barré syndrome), spina bifida, and chronic or degenerative neurological problem.

All study procedures of the UK Biobank were approved by the North-West Multi-Centre Research Ethics Committee. All participants provided informed consent to participate in the study. Details on the UK Biobank Ethics and Governance framework are provided at https://www.ukbiobank.ac.uk/media/0xsbmfmw/egf.pdf.

### UK Biobank: Brain imaging acquisition

The UK Biobank brain imaging data was acquired using 3.0 Tesla Siemens Skyra scanner with a standard Siemens 32–channel head coil—across three imaging centers with identical scanners. The imaging acquisition parameters are described in detail at https://biobank.ctsu.ox.ac.uk/crystal/refer. cgi?id=2367 and brain imaging documentation is provided at https://biobank.ctsu.ox.ac.uk/crystal/refer.cgi?id=1977 (also see Miller et al. [138]).

T1–weighted structural data was acquired using a 3D MPRAGE sequence with the following parameters: repetition time (TR) = 2 000 ms, inversion time (TI) = 880 ms, echo times (TE) = 2.01 ms, spatial resolution = 1 × 1 × 1 mm^3^, number of echoes = 4, flip angle = 8° and field of view of 208 × 256 × 256 mm. T2–weighted fluid-attenuated inversion-recovery (FLAIR) structural data was acquired using a 3D SPACE sequence. Acquisition parameters for the T2–weighted sequence were: TR = 5 000 ms, TE = 395 ms, spatial resolution = 1.05 *×* 1 *×* 1 mm^3^, turbo factor = 284, and field of view of 192 *×* 256 *×* 256 mm.

Diffusion data was acquired using an EPI sequence with the following parameters: TR = 3 600 ms, flip angle = 78°, 50 distinct directions on 2 shells, *b* = 1 000*/*2 000 s*/*mm^2^, along with 10 *b* = 0 s*/*mm^2^ images, and field of view of 104 × 104 mm, imaging matrix of 52 × 52, 72 slices with thickness of 2 mm.

ASL data was acquired using pCASL and a 2D multi-band EPI sequence. Pseudo-continuous ASL data was acquired with labeling duration of 1 800 ms and five post-labeling delays of 400 ms, 800 ms, 1 200 ms, 1 600 ms, and 2 000 ms. One label/control pair per post-labeling delay is acquired. To calibrate perfusion measurements into units of ml*/*100g*/*min, one M0 calibration image (effective TR = 5 s) without labeling was also acquired. The spatial resolution for this imaging modality was 3.4 × 3.4 × 4.5 mm^3^. Summary measures for blood perfusion and ATT were provided for major brain lobes and subcortical structures. We included 50 ASL-based imaging-derived phenotypes (IDPs) per participant.

### UK Biobank: Brain imaging measurements

To relate brain measurements to peripheral biomarkers, we incorporated imaging-derived phenotypes (IDPs) provided by the UK Biobank repository. All IDPs were obtained using the standardized UK Biobank processing pipelines, available at: https://git.fmrib.ox.ac.uk/falmagro/UK_biobank_pipeline_v_1. Specifically, we used diffusion tensor imaging measures (FA, MD), microstructural measures (ISOVF, ICVF), in addition to regional volumes, cortical thickness, cortical surface areas, white matter hyperintensity (WMH), blood perfusion and ATT measures.

Cortical morphometric features—including thickness, surface area, and volume—were extracted using the Desikan–Killiany (DK) atlas (62 parcels). Diffusion-based metrics—including FA, MD, ICVF, and ISOVF—were parcellated using the ICBM-DTI81 white-matter labels atlas (48 white matter tract labels). We further incorporated 25 regionally defined perfusion- and ATT-related measures. These included values for the right and the left hemispheres’ frontal lobe, occipital lobe, parietal lobe, temporal lobe, cerebellum, caudate, putamen, thalamus, cerebrum white matter and internal carotid artery vascular territory in gray matter. We also used IDPs for mean blood perfusion and ATT in vertebrobasilar arteries vascular territories in gray matter, cerebral white matter with >50% cerebral partial volume, cortical gray matter and whole-brain gray and white matter.

Subcortical structures were defined using Freesurfer segmentation (aseg), from which volumetric measures were extracted for the following bilateral regions: accumbens area, thalamus proper, pallidum, putamen, hippocampus, caudate, amygdala, choroid plexus, cerebral white matter, cerebellar white matter, cerebellar cortex, cerebral cortex, and ventral diencephalon. In addition, whole-brain summary measures were included, such as brain stem, BrainSeg, BrainSegNotVent, BrainSegNotVentSurf, total gray matter, and subcortical gray matter volumes. We also included subcortical volumes derived from FSL-FIRST for key regions including the accumbens, thalamus, pallidum, putamen, hippocampus, caudate and amygdala. Regional gray matter volumes were obtained using FSL-FAST for several subcortical areas, including the thalamus, ventral striatum, pallidum, putamen, hippocampus, caudate, and amygdala. Finally, two imaging-derived phenotypes (IDPs) specifically characterizing white matter hyperintensity (WMH) burden were included: total WMH volume estimated from T1– and T2–FLAIR images, and the total volume of white matter hypointensities across the whole brain.

Altogether, the brain feature set included 490 values. We ensured that no subject included in the multi-model mapping had more than 40 missing values for brain measures; any subject exceeding this threshold was excluded from the analysis.

### UK Biobank: Biological samples and vital signs

The UK Biobank dataset included a comprehensive array of participant-specific blood and urine laboratory results in addition to obesity-related measures, blood pressure and age. The laboratory-based assessments included C-reactive protein, total protein, testosterone, glucose, HbA1c, cholesterol, HDL, LDL, triglycerides, urea, calcium, bilirubin, creatinine, vitamin D, AST, ALT, ALK, sodium urine, potassium urine, and systolic and diastolic blood pressures. Furthermore, participants’ directory included data on biomarkers as age, BMI, body fat percentage, waist and hip circumference.

UK Biobank is a longitudinal study, laboratory results were acquired only at the initial assessment visit, while blood pressure and obesity-related measures (e.g., BMI, hip and waist circumference and body fat percentage) were acquired at both the initial assessment and the follow-up brain imaging visits. We included age at baseline and age at imaging visit in the original PLS model. We also performed an additional PLS analysis in which the effect of age effect was regressed out from all brain imaging and physiological features.

Among the 1 582 female and 1 431 male participants, there were some features with missing values. These were substituted with the median of the sex-specific corresponding columns. For information on number of missing values per biomarker measure see Table. S2).

### UK Biobank: Fluid intelligence assessment

All UK Biobank participants completed a fluid intelligence assessment at the time of imaging. The assessment comprised 13 multiple-choice questions administered within a two-minute time frame, and was designed to quantify problem-solving and reasoning abilities in participants. Questions were presented individually on screen with 3–5 response options. For each question, participants could select their chosen answer, or “Do not know”, and “Prefer not to answer” alternatives. The assessment evaluated verbal reasoning (e.g., analogical relationships: “Bud is to flower as child is to…?” with options including Grow, Develop, Improve, Adult, Old) and numerical reasoning (e.g., sequence completion: “150…137…125…114…104… What comes next?” with options 96, 95, 94, 93, 92). The fluid intelligence score was calculated as the total number of correct responses within the two-minute limit [55].

When calculating correlations between PLS scores and fluid intelligence scores, data of individuals with missing fluid intelligence scores were excluded. Missing scores totaled 159 among females, and 121 among males.

### Atlases

Cortical features in the HCP–Aging dataset were extracted using the Schaefer-400 functional atlas [182]. White matter features were extracted using the Johns Hopkins University (JHU) white-matter tractography atlas (threshold 25%) [89, 226]. WMH values were obtained using the Hammer atlas available at https://zenodo.org/records/7930159.

In the UK Biobank dataset, cortical thickness, and surface area were provided for each parcel of the Desikan-Killiany-Tourville (DKT) atlas. This information was derived using the *FreeSurfer* v6.0.0. For cortical blood perfusion and arterial transit time, broader anatomical labels (frontal lobe, insula, occipital cortex, parietal lobe, and temporal lobe) were used due to the low spatial resolution of the ASL imaging.

Regional subcortical volumes in the UK Biobank were obtained with the Functional Magnetic Resonance Imaging of the Brain (FMRIB)’s Integrated Registration and Segmentation Tool (FIRST). For subcortical cerebral blood perfusion and arterial transit time, only major structures such as caudate, cerebellum, and thalamus were included.

In UK Biobank data, FA, MD, ICVF, and ISOVF measures were summarized across 48 standard parcels of the JHU atlas.

### Cortical thickness

Cortical thickness quantifies the width of cortical gray matter. The individual-level cortical thickness maps in HCP–A dataset were derived through the HCP Minimal Preprocessing Pipeline [67]. In short, this pipeline used both participant-specific bias-corrected T1–/T2–weighted structural data to segment the cortical gray and white matter and performed a surface reconstruction using FreeSurfer. Next, cortical thickness was estimated as the geometric distance between the white and pial surfaces.

The individual-level cortical thickness maps in UK Biobank were also constructed by incorporating both T1– and T2–weighted structural data using the *FreeSurfer* pipeline.

### Cortical Myelin

T1/T2 ratio quantifies the cortical myelin profile. T1/T2 ratio is affected by the microstructural characteristics of cortical gray matter. Division of the T1– to T2–weighted image enhances myelin contrast to noise ratio [68, 201]. Specifically, we used the RF transmit (B1+) field corrected myelin maps following recommendation from Glasser et al. [66]. Myelin was only assessed in HCP–A dataset.

### Fractional anisotropy and mean diffusivity

Fractional anisotropy (FA) quantifies the anisotropy of water diffusion, and ranges from 0 to 1. FA = 1 indicates that within the region of interest, all water movements occur in the same direction, while FA = 0 indicates that there is no dominant directionality in the movements of water molecules. In other words, FA is a measure that examines the “relative” directional water diffusion along the principal axis of diffusion versus diffusion in other direction. Higher FA may indicate greater fiber density, lower membrane permeability or greater myelin. Higher FA can also be a result of disproportionate decrease in one or more of the non-dominant fiber bundles. Notably, constant FA of a region is not indicative of unchanged underlying tissue; in a case where bundles within a region are all damaged in the same way, FA—which is a relative measure of water diffusivity—stays constant [58].

Mean diffusivity (MD) quantifies the overall anisotropy capability of water within a region of interest (mean of diffusion along the three tensor dimensions). MD quantification is more robust and interpretable (than FA) in the presence of crossing fibers, and can assess how the tissue is constraining the water diffusion in a region of interest [58].

Both FA and MD measures were derived from diffusion tensor imaging. Cortical FA and MD were only assessed in HCP–A dataset, while white matter FA and MD were assessed in both datasets.

### Functional connectivity strength

Functional connectivity (FC) quantifies the synchronization of fluctuation in BOLD signals across brain regions. Functional MRI data was preprocessed using the HCP Minimal Preprocessing Pipeline. For detailed preprocessing steps, refer to the cited reference [67]. The vertex-wise functional MRI timeseries were initially demeaned and subsequently parcellated using the Schaefer-400 functional atlas [182]. The parcellated time-series were then *z*scored and concatenated across participant runs. Each participant’s unified time-series was used to derive functional connectivity matrix. The functional connectome was computed by calculating the Pearson correlation coefficient between pairs of regional time-series. To construct the functional connectivity strength map, we computed the absolute value of all functional connectivity edges and sum the edges connected to each region for each participant (an array of size 400 per participant). FC strength was only assessed in HCP–A dataset.

### Structural connectivity strength

Structural connectivity (SC) quantifies the density of white matter tracts across brain regions. Diffusion MRI data was preprocessed using the HCP Minimal Preprocessing Pipeline. We next used *probtrackx* to build the participant specific connectomes using the schaefer-400 cortical parcellation. We used the default settings when running the *probtrackx*. To construct the structural connectivity strength map, we computed the absolute value of all structural connectivity edges and summed the edges connected to each region for each participant (an array of size 400 per participant). SC strength was only assessed in HCP–A dataset.

### White matter hyperintensity

White matter hyperintensities (WMHs) are regions detectable on structural brain MRI, appearing hyperintense on T2–weighted and fluid-attenuated inversion-recovery (FLAIR) images, and hypointense on T1–weighted images. They are associated with microangiopathy and chronic ischemia. Histopathological studies indicate that WMHs reflect tissue damage ranging from slight disentanglement of the matrix to varying degrees of myelin and axonal loss [166].

In HCP–A dataset, BISON (Brain tISsue segmentatiON) automatic segmentation tool was used to segment WMHs [43]. BISON combined a random forest classifier with a collection of location and intensity features obtained from a library of manually segmented scans to detect WMHs. T2–weighted and T1–weighted images were utilized for WMH segmentation in HCP–A. The WMH segmentations were visually quality controlled by an experienced rater (R.M.). The number of voxels designated as WMH (measured in mm^3^ in the standard space (i.e., adjusted for intracranial volume) in each brain lobe and hemisphere (based on Hammer’s lobar atlas) as well as across the entire brain were used to define regional and global WMH volumes, respectively. WMH volumes were log-transformed to obtain a normal distribution.

In UK Biobank, we used two IDPs to quantify ischemia-related structural changes: total volume of white matter hyperintensities from T2 FLAIR and T1 images [74], and volume of white matter hypointensities in the whole brain, generated based on T1 images.

### Intracellular volume fraction and isotropic volume fraction

Intracellular volume fraction (ICVF) quantifies the density of neurites (axons and dendrites). Isotropic volume fraction (ISOVF) quantifies the proportion of free water (extracellular water diffusion). Higher ICVF and lower ISOVF indicate better tissue microstructural organization. These measures were calculated by feeding the diffusion MRI data in the Neurite Orientation Dispersion and Density Imaging (NODDI) modeling using the Accelerated Microstructure Imaging via Convex Optimization (AMICO) tool [44, 243]. ICVF and ISOVF were only assessed in the UK Biobank dataset.

### Partial least squares

Partial least squares (PLS) analysis was performed using the *pyls* toolbox (https://github.com/netneurolab/pypyls/). PLS analysis is a technique used to capture the multivariate relationship between two data matrices (not necessarily the causal effect between the datasets) [110, 132, 133]. In this report, the investigated data matrices were “brain imaging feature maps” (*N*_participants_ × *N*_imaging features_) and peripheral biomarker (*N*_participants_ × *N*_biomarkers_). Each column in each data matrix was first *z*-scored; next, the covariance (correlation) between normalized brain imaging features (**X**) and biomarkers (**Y**) was computed. The covariance matrix was decomposed (**X**^*′*^**Y**) using the singular value decomposition (SVD):

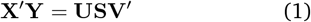

here **U** and **V** were orthonormal matrices of left and right singular vectors and **S** was a diagonal matrix of singular values. Each triplet of a left singular vector, a right singular vector, and a singular value constituted a latent variable. Singular vectors weighted the contribution of original features (regional brain imaging features and biomarkers) to the overall multivariate pattern. To quantify the extent to which individual participants expressed the multivariate pattern captured by a latent variable, participant-specific brain imaging and biomarker scores were calculated. Scores were computed by projecting the original data onto the respective singular vector weights, such that each individual was assigned a brain imaging score and a biomarker score:

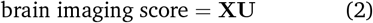

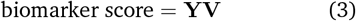

Next, PLS loadings of a latent variable were computed by calculating the Pearson’s correlation between participant-specific biomarker measures (or brain imaging measures) and the respective score pattern. The proportion of covariance explained by each latent variable was quantified as the ratio of the squared singular value to the sum of all squared singular values. The statistical significance of each latent variable was determined by permutation test. The testing process involved randomly permuting the order of observations (i.e., rows) of data matrix **X** for a total of 1 000 repetitions, followed by constructing a set of “null” brain-biomarker correlation matrices. These “null” correlation matrices were then subjected to SVD, to generate a distribution of singular values under the null hypothesis that there was no association between brain imaging features’ pattern and participants’ biomarkers. A non-parametric *p*-value could be estimated for each given latent variable as the probability that a permuted singular value exceeds the original, nonpermuted singular value.

The reliability of individual biomarker contribution to the latent variable was evaluated using bootstrap resampling. Participants (rows of data matrices **X** and **Y** were randomly sampled with replacement across 1 000 repetitions, resulting in a new set of correlation matrices that were subsequently subjected to SVD. This procedure generated a sampling distribution for each individual weight in the singular vectors. For each biomarker, a bootstrap ratio was computed as the ratio of its singular vector weight to its bootstrap-estimated standard error. High bootstrap ratios indicated biomarkers that significantly contribute to the latent variable and are stable across participants.

Finally, we used cross-validation to assess the generalizability of PLS results [81, 140]. We assessed the out-of-sample correlation between brain imaging features and biomarker scores. We use 100 randomized train-test splits of the dataset; in each random split, we used 80% of the data for training and 20% of the data for testing. In each iteration, PLS was applied to the training data (**X**_train_ and **Y**_train_) to estimate singular vector weights (**U**_train_ and **V**_train_); test data was then projected onto these derived weights to compute participant-specific scores (**X**_test_*U*_train_ and **Y**_test_*V*_train_). The correlation between brain and biomarker scores was evaluated for the test sample. This procedure led to 100 correlation values and established a distribution of outof-sample correlation values. To assess the statistical significance of these out-of-sample correlation values, we conducted permutation tests (100 repetitions). During each permutation, we shuffled the matrix rows and repeat the analysis to create a null distribution of correlation coefficients between brain imaging and physiological scores in the test sample. This null distribution was then used to estimate a non-parametric *p*-value, by calculating the proportion of null correlation coefficients that were greater than or equal to the mean original out-of-sample correlation coefficient.

### Generalized additive model for location, scale, and shape

Generalized additive model for location, scale, and shape (GAMLSS) was implemented using the *gamlss* R package available at https://github.com/gamlss-dev/ [198]. We used sex-stratified GAMLSS modeling approach to regress-out the non-linear effect of age from features including, brain measurements, peripheral biomarkers and cognitive scores. In this framework, each feature of interest is assumed as a random variable (*Y*) following the normal distribution (𝒩):

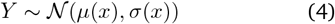

here, 𝒩 distribution parameters—mean (*µ*), standard deviation (*σ*)—can be modeled are modeled as the fractional polynomial functions of the explanatory variable(age; *x*). We used the fp() function to determine the best-fitting two-term fractional polynomials from the predefined set of power values: − 2, − 1, − 0.5, 0, 0.5, 1, 2, 3. The optimal set of powers is chosen automatically through iterative model fitting to best capture non-linear relationships between age and distribution parameters.

For the UK Biobank dataset, we regressed out the effects of both age and imaging center by using nonlinear models that included center ID as a fixed covariate. This ensured that remaining variation in the features reflected neither age nor center-related biases.

### Statistics and null models

To assess the effect of spatial auto-correlation on spatial associations between two cortical brain maps, we used the so-called spatial auto-correlation preserving permutation tests, commonly referred to as “spin tests” [221]. Briefly, brain phenotypes were projected to spherical projection of the fsaverage surface. This involved selecting the coordinates of the vertex closest to the center of mass for each parcel. These parcel coordinates were then randomly rotated, and original parcels were reassigned to the value of the closest rotated parcel (*N* repetitions; throughout the manuscript *N* = 1 000). For parcels where the medial wall was the closest, we assigned the value of the next closest parcel instead. Following these steps, we obtained a series of randomized brain maps that have the same values and spatial auto-correlation as the original map but where the relationship between values and their spatial location had been permuted. These maps were then used to generate null distributions of desired statistics. Throughout the manuscript, whenever the spatial correspondence between two brain maps was tested for, “spin test” was carried out for maps parcellated with the Schaefer-400 cortical atlas [182]. Notably, this parcellation was chosen because it divides the cortex into relatively homogeneous parcel sizes, and its number of parcels was comparable to the estimated number of distinct human neocortical areas [151, 220].

To assess the statistical significance of the association between cognitive scores and brain loadings, we performed permutation testing. We first calculated Spearman’s rank correlation between the observed cognitive scores and brain loadings. Next, we permuted the cognitive scores 1 000 times and recalculated the correlation for each permutation to generate a null distribution. Two-sided, non-parametric p-values were then derived by comparing the observed correlation against this null distribution.

Furthermore, significance of PLS latent variables was assessed by permutation tests. The generalization of PLS results was assessed using crossvalidation approach; see *Methods: Partial least squares* for more details. Note that throughout the manuscript statistical significance was assumed at pvalue smaller than 0.05.

## Data and code availability

The Human Connectome Project-Aging (HCP–A) is accessible through https://www.humanconnectome.org/study/hcp-lifespan-aging/ [22, 82, 194]. The UK Biobank data is accessible through https://www.ukbiobank.ac.uk/.

All code used to perform the analyses can be found on GitHub at https://github.com/netneurolab/Farahani_Metabolic_Health/. The code directly relies on open-source Python packages including, including NumPy (version 1.21.6) [83, 219], SciPy (version 1.7.3) [224], pandas (version 1.3.5) [134], seaborn (version 0.12.2) [230], Matplotlib (version 3.5.3) [91], statsmodels (version 0.13.5) [185], bctpy (version 0.6.1) [178], Nilearn (version 0.10.1, see [2]), NiBabel (version 4.0.2) [24], netneurotools (version 0.2.3) [126], and rpy2 (version 3.5.17) [64].

Parcellation atlases, including the Schaefer400 functional atlas [182], and Johns Hopkins University (JHU) white matter tractography atlas [89, 226], can be obtained from https://github.com/ThomasYeoLab/CBIG/tree/master/stable_projects/brain_parcellation/Schaefer2018_LocalGlobal/Parcellations, and https://web.mit.edu/fsl_v5.0.10/fsl/doc/wiki/Atlases.html, respectively. The GAMLSS models are implemented using GAMLSS R package (version 5.4-22) available at https://github.com/gamlss-dev/ [198]. The PLS analysis is performed using the pyls toolbox (version 0.0.1) available at https://github.com/netneurolab/pypyls/. All brain plots in the manuscript are visualized using Connectome Workbench (version 1.5.0), available at https://www.humanconnectome.org/software/get-connectome-workbench/ [125].

## Competing interests

The authors declare no competing interests.

## Acknowledgments

We thank all participants and staff of the HCP–A and UK Biobank datasets. We thank Justine Hansen, Vincent Bazinet, Filip Milisav, Andrea Luppi, Yigu Zhou, Tahmineh Taheri, and Moohebat Pourmajidian for their comments and suggestions on the manuscript. This research has been conducted using the UK Biobank Resource under Application Number 45551. AF acknowledges support from the Molson Foundation. BM acknowledges support from the NSERC, Canadian Institutes of Health Research (CIHR), Brain Canada Foundation Future Leaders Fund, the Canada Research Chairs Program, the Michael J. Fox Foundation and the Healthy Brains for Healthy Lives initiative. The funders have no role in study design, data collection and analysis, decision to publish or preparation of the paper.

---

**Figure S1.**
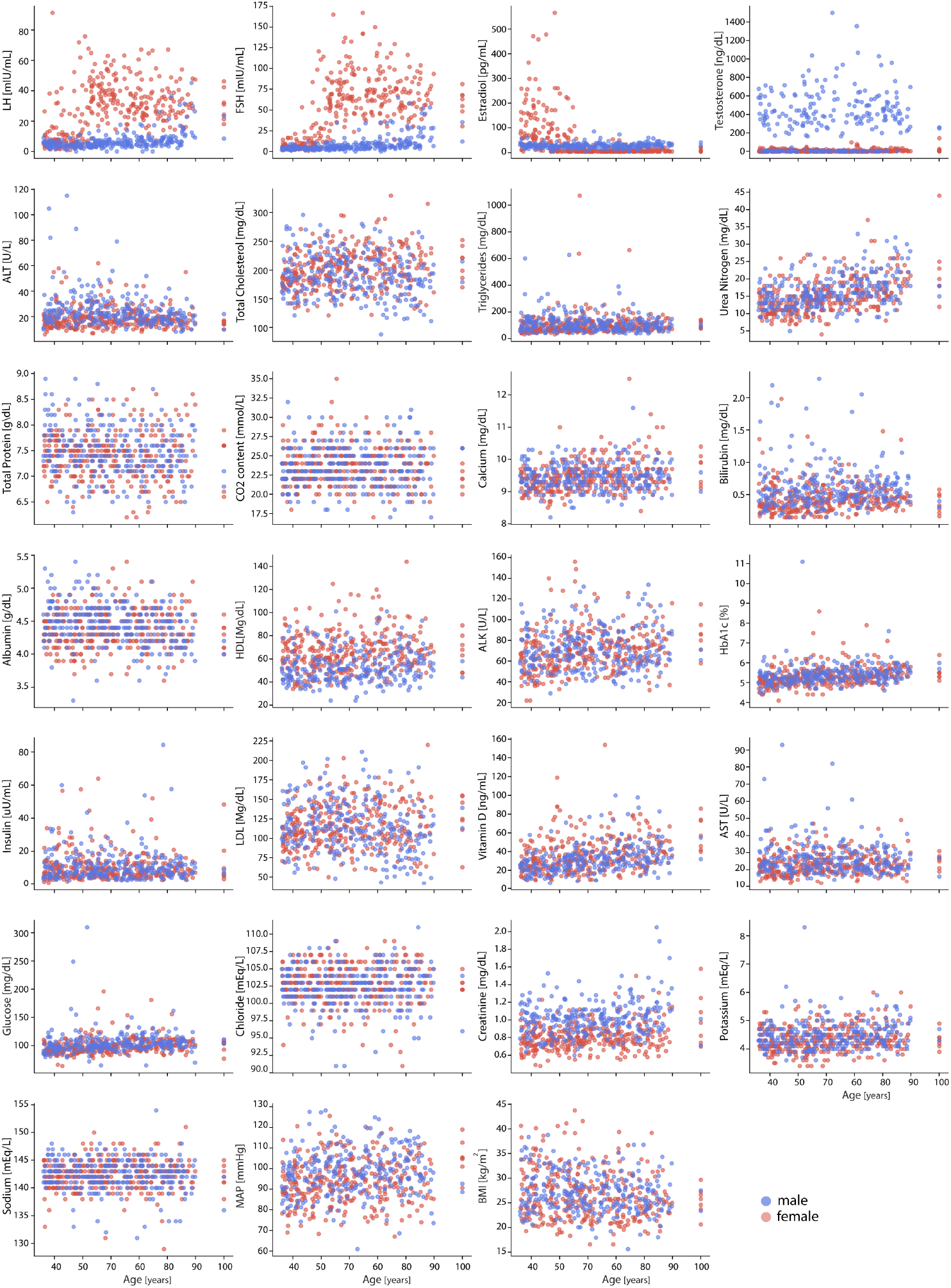
HCP–A biomarkers versus age. Relationship between participants’ age (x-axis) and raw biomarker values (y-axis) (blue: male, red: female).

**Figure S2.**
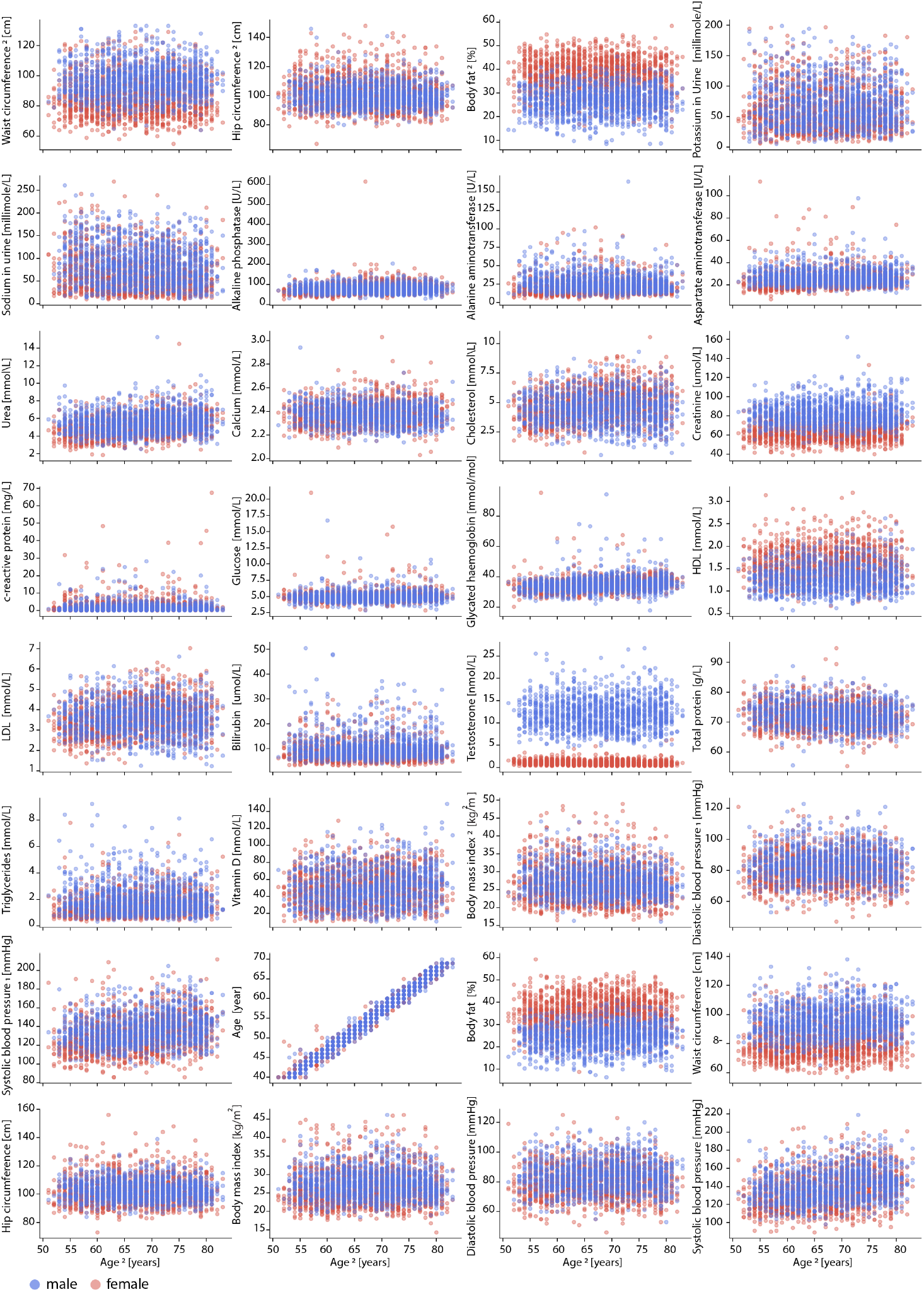
UK Biobank biomarkers versus age. Relationship between participants’ age at the time of brain imaging (x-axis) and raw biomarker values (y-axis) (blue: male, red: female). For biomarkers measured at both initial assessment and follow-up imaging visits, values corresponding to the imaging visit are denoted with the subscript “2”.

**Figure S3.**
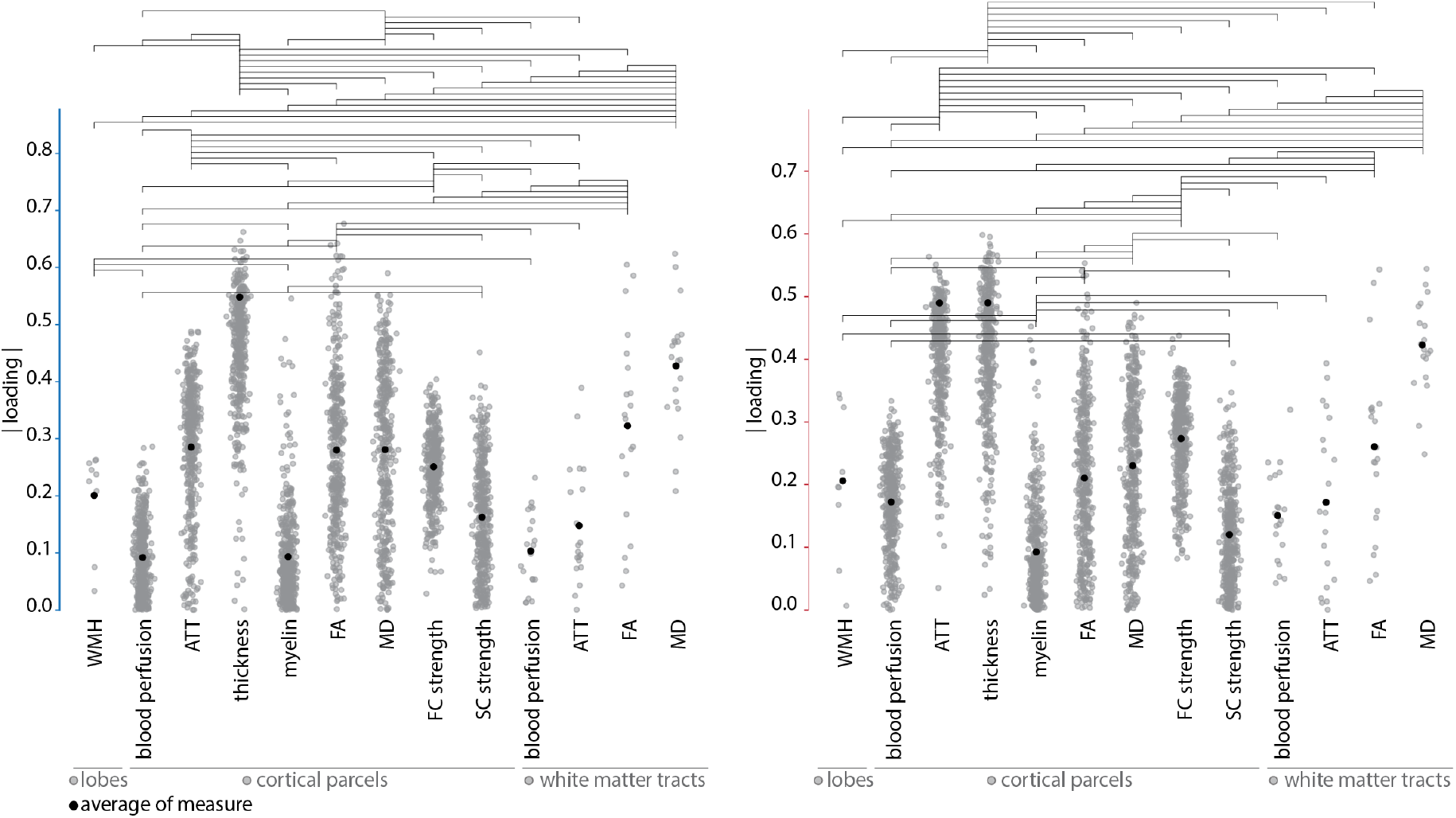
Statistical comparison of absolute brain loading values for the first latent variable. Horizontal lines indicate statistical significance (*p <* 0.05, FDR corrected). Black dots represent mean absolute loading values. Also see Fig. 1b.

**Figure S4.**
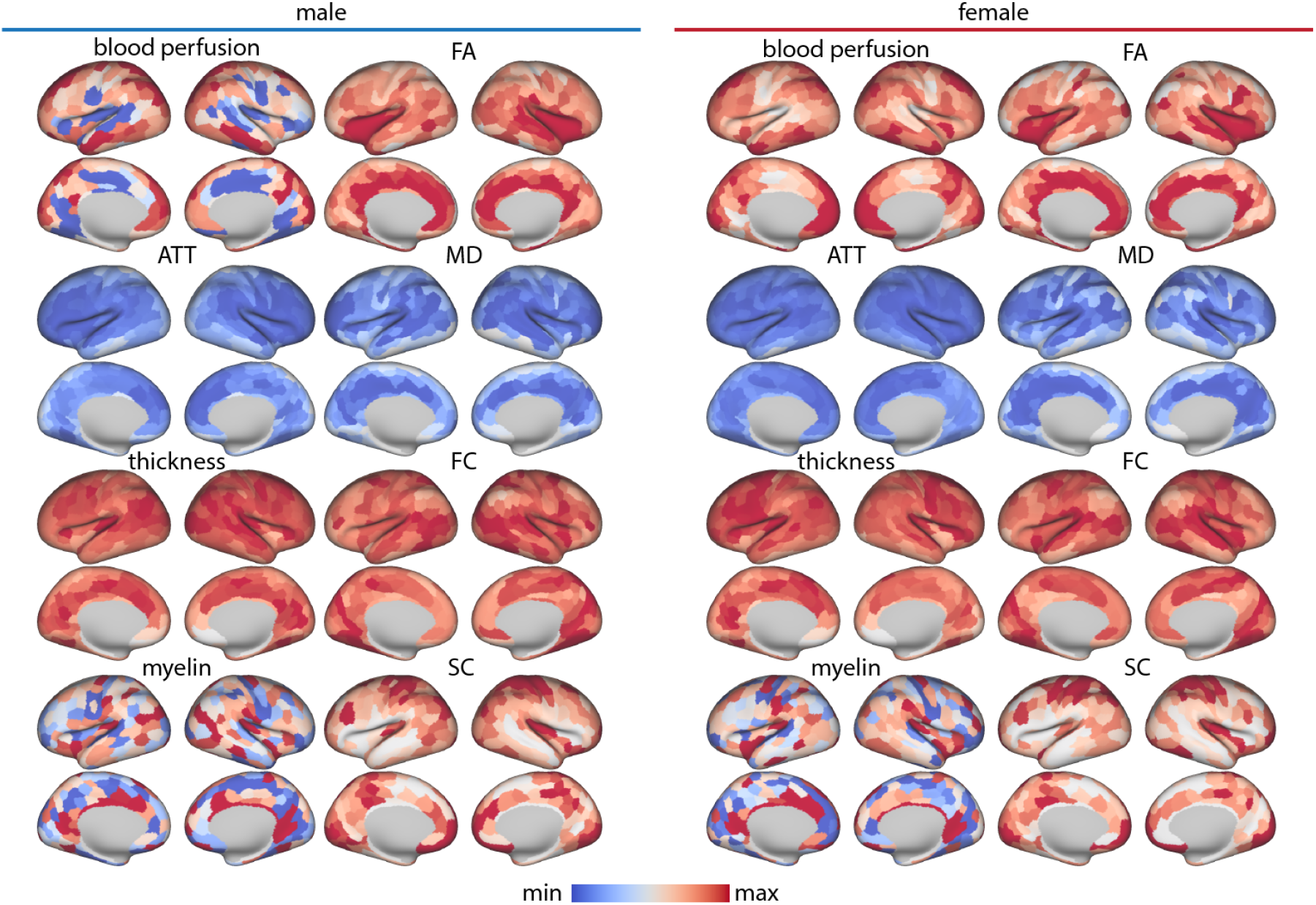
LV–I Brain loadings. Brain loadings are shown on both right and left hemispheres. Loadings are shown on the fsLR inflated cortical surfaces.

**Figure S5.**
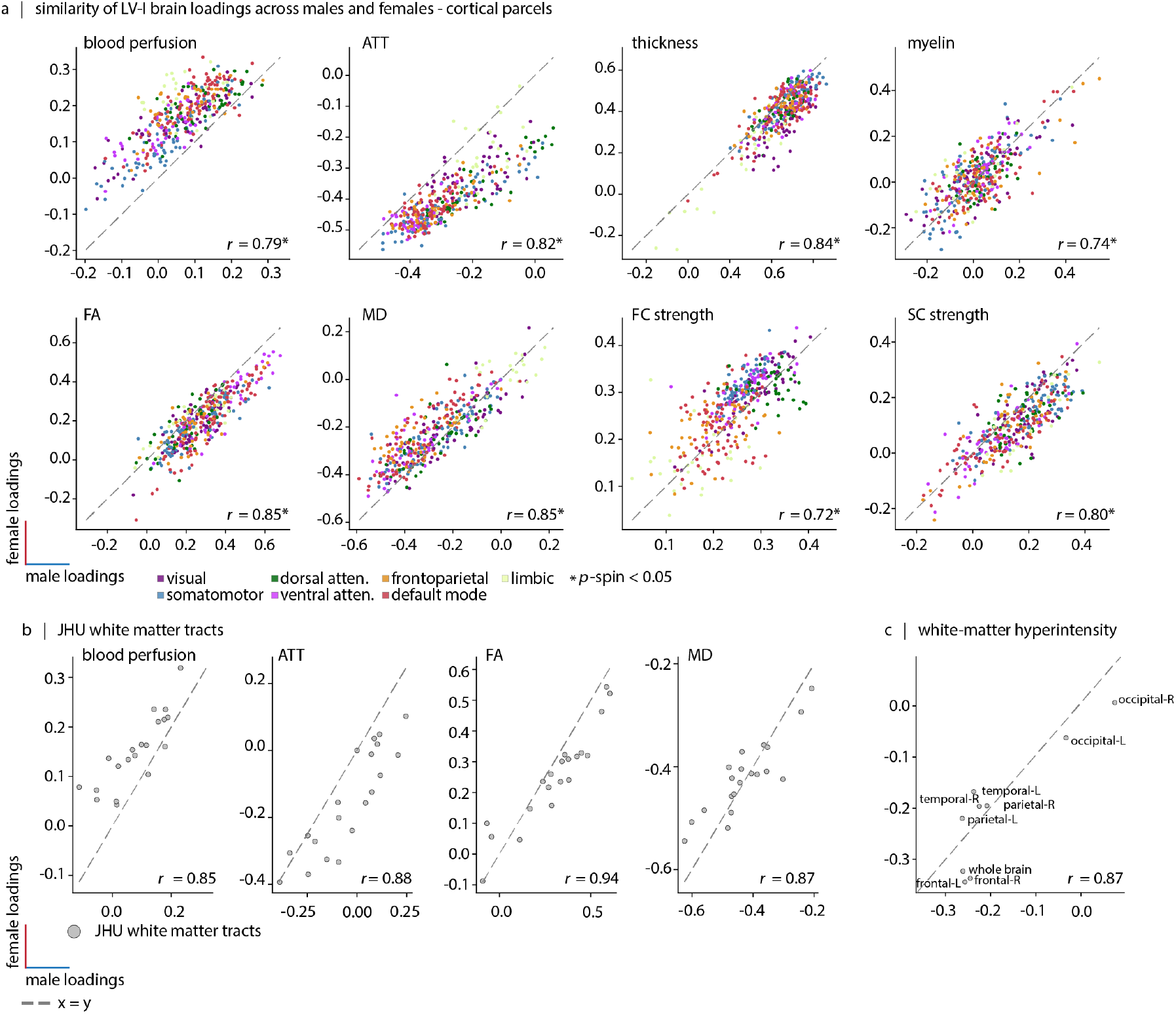
Similarity of first latent variable brain loadings between males and females. (a) Cortical brain loadings comparison. In the scatter plots, each dot represents a brain region defined by the Schaefer-400 parcellation [182]; the dots are color-coded based on the Yeo-7 functional resting-state networks [208]. The *x*-axis shows male brain loadings and the *y*-axis shows female brain loadings. (b) White matter tract loadings comparison. The scatter plots show the similarity of brain loadings between males (*x*-axis) and females (*y*-axis) for white matter tract measurements including blood perfusion, arterial transit time (ATT), fractional anisotropy (FA) and mean diffusivity (MD). (c) White matter hyperintensity loadings comparison. The scatter plots show the similarity of brain loadings between males (*x*-axis) and females (*y*-axis) for white matter hyperintensity measurements.

**Figure S6.**
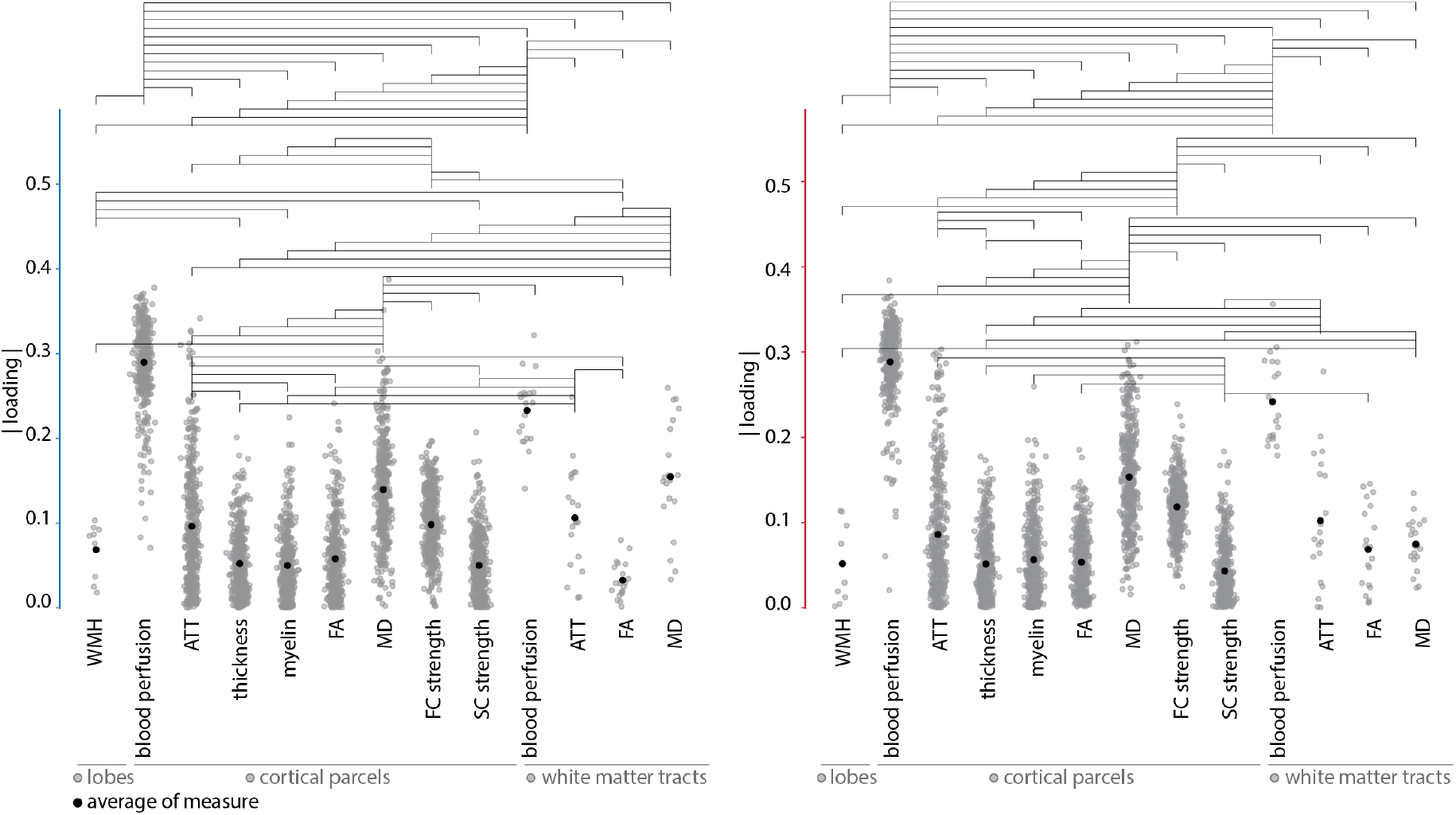
Statistical comparison of absolute brain loading values for the second latent variable. Blood perfusion loadings have the greatest absolute values among all brain measures examined. Horizontal lines indicate statistical significance (*p <* 0.05, FDR corrected). Black dots represent mean absolute loading values. Also see Fig. 3b.

**Figure S7.**
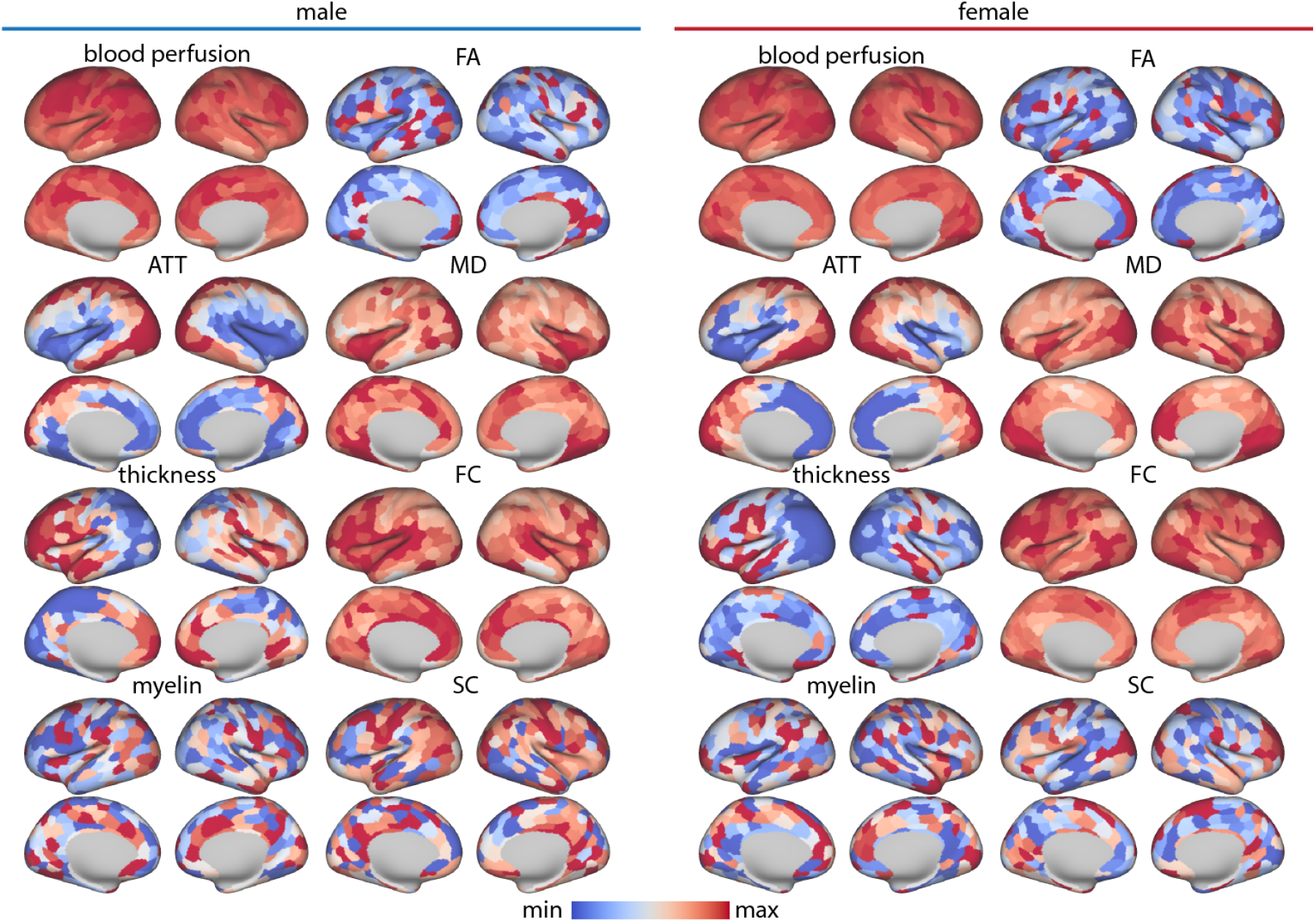
LV–II Brain loadings. Brain loadings are shown on both right and left hemispheres. Loadings are shown on the fsLR inflated cortical surfaces.

**Figure S8.**
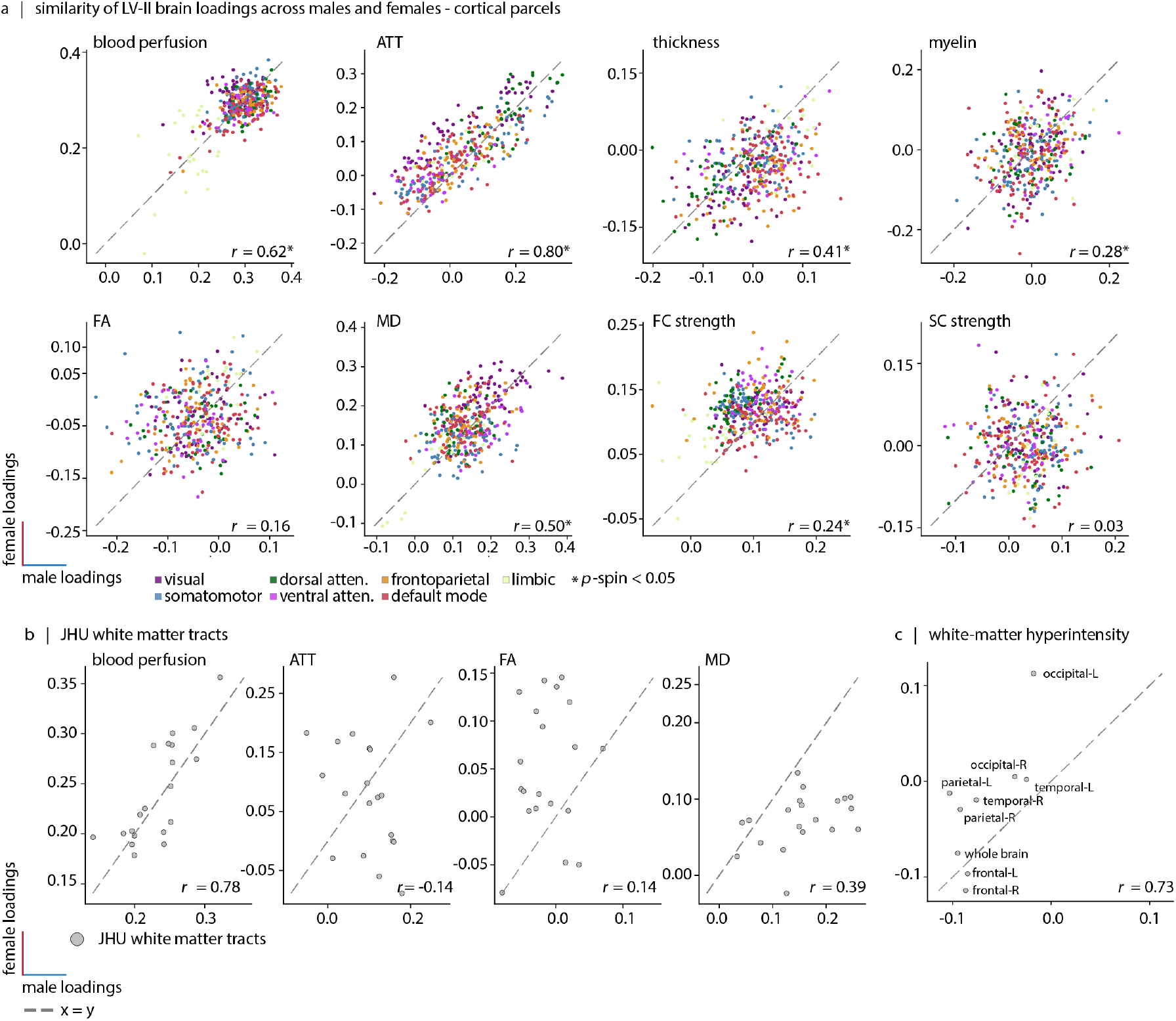
Similarity of second latent variable brain loadings between males and females. (a) Cortical brain loadings comparison. In the scatter plots, each dot represents a brain region defined by the Schaefer-400 parcellation [182]; the dots are color-coded based on the Yeo-7 functional resting-state networks [208]. The *x*-axis shows male brain loadings and the *y*-axis shows female brain loadings. (b) White matter tract loadings comparison. The scatter plots show the similarity of brain loadings between males (*x*-axis) and females (*y*-axis) for white matter tract measurements including blood perfusion, arterial transit time (ATT), fractional anisotropy (FA) and mean diffusivity (MD). (c) White matter hyperintensity loadings comparison. The scatter plots show the similarity of brain loadings between males (*x*-axis) and females (*y*-axis) for white matter hyperintensity measurements.

**Figure S9.**
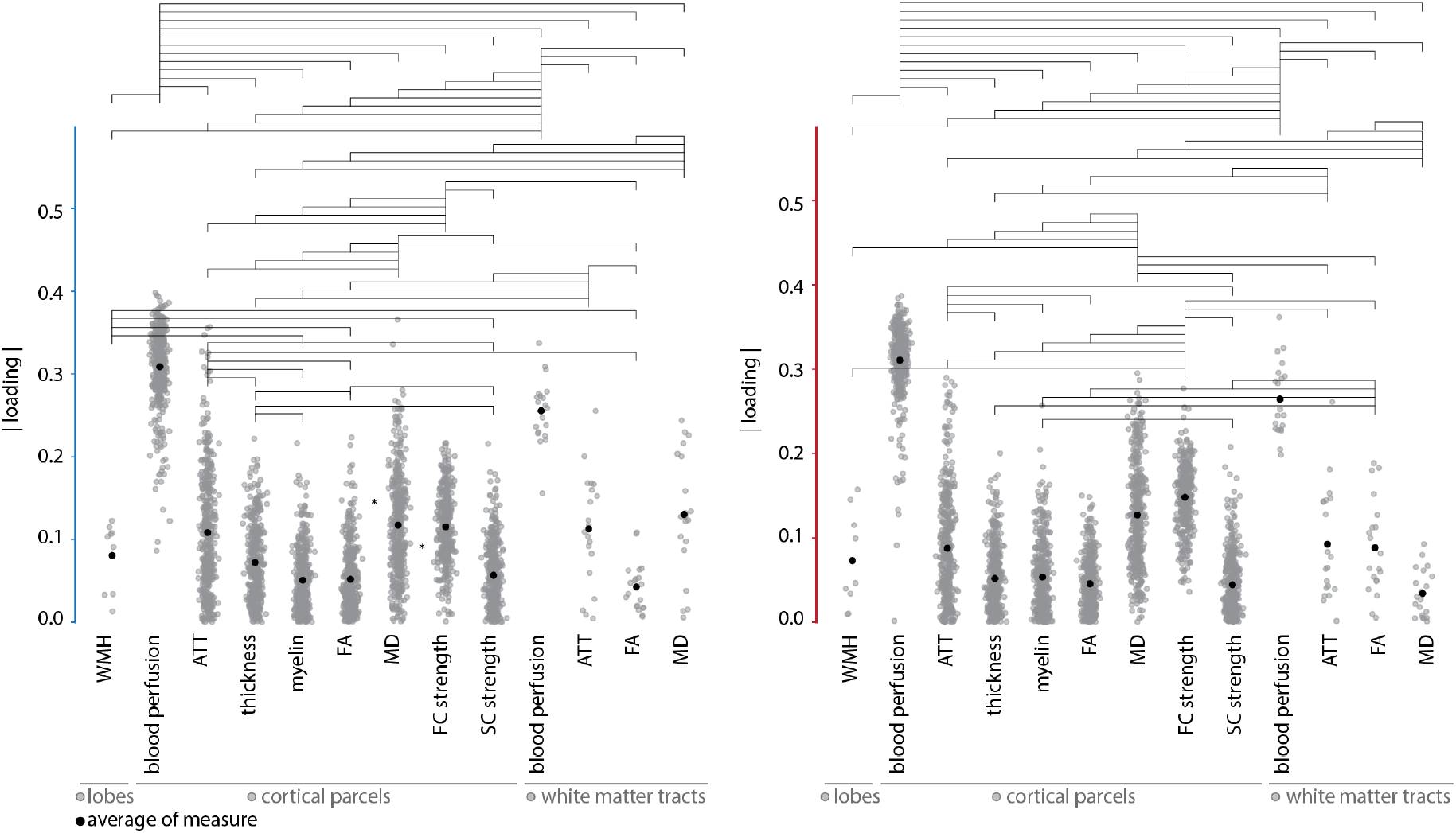
Statistical comparison of absolute brain loading values for the first latent variable - no age effect. Blood perfusion loadings have the greatest absolute values among all brain measures examined. Horizontal lines indicate statistical significance (*p <* 0.05, FDR corrected). Black dots represent mean absolute loading values. Also see Fig. 5b.

**Figure S10.**
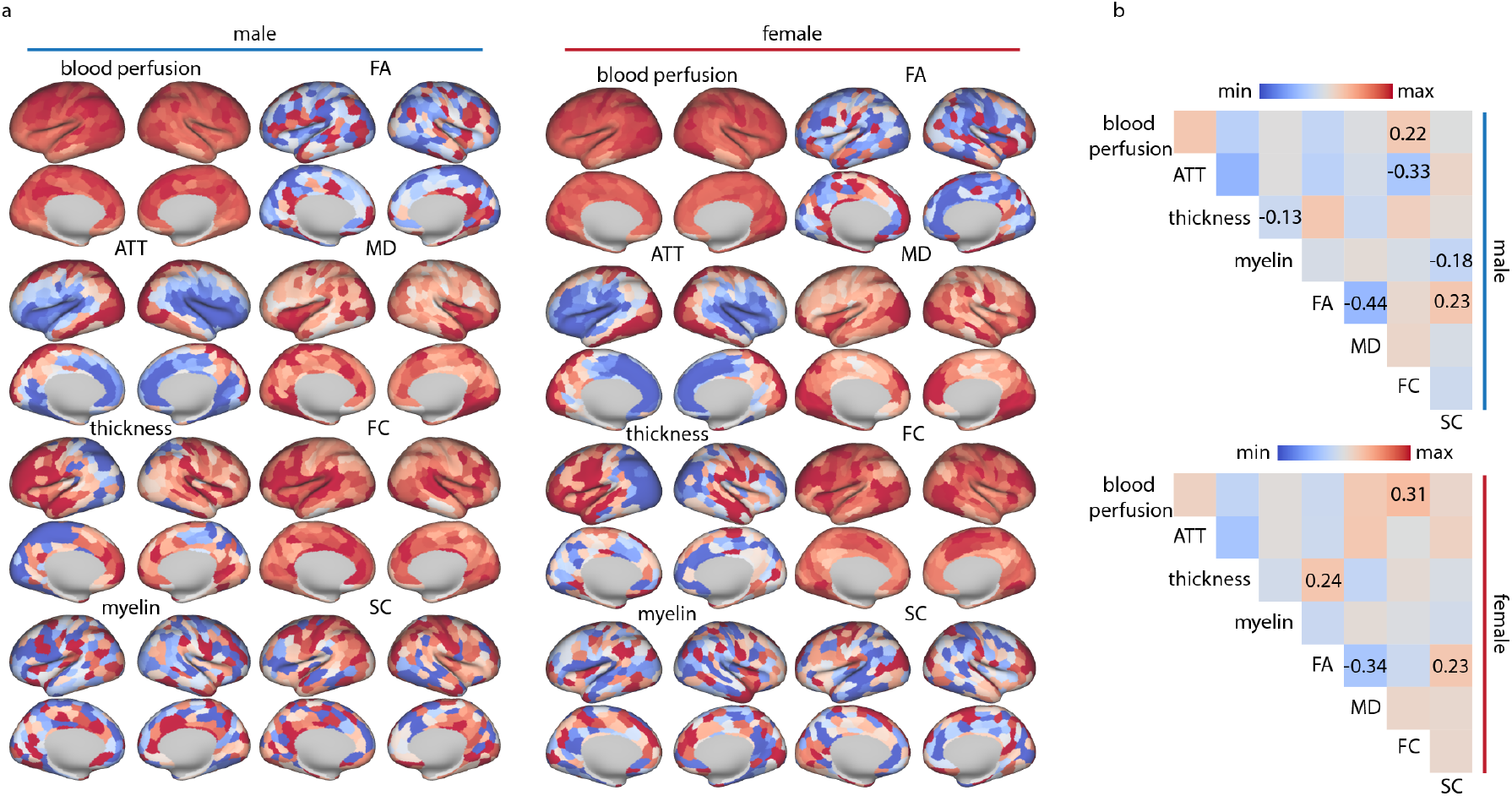
LV–I^*′*^ brain loadings - no age effect. (a) Brain loadings for each included brain feature. This indicates that participants who more strongly express the biomarkers’ pattern illustrated in Fig. 5a (e.g., those who lower BMI), have higher blood perfusion. Brain maps are shown on the fsLR inflated cortical surfaces. (b) Similarity of brain loadings across included brain features. Associations that remain significant after controlling for spatial autocorrelation and false discovery rate (FDR) correction are marked with their corresponding correlation values (FC and blood perfusion: *p*_spin_ = 0.042 (males), *p*_spin_ = 1.86 *×* 10^−2^ (females); FA and MD: *p*_spin_ = 6.99 *×* 10^−3^ (males), *p* = 0.014 (females); FA and SC: *p*_spin_ = 6.99 *×* 10^−3^ (males), *p*_spin_ = 0.014 (females); myelin and SC: *p*_spin_ = 6.99 *×* 10^−3^ (males); ATT and FC: *p*_spin_ = 6.99 *×* 10^−3^ (males); thickness and myelin: *p*_spin_ = 0.042 (males); thickness and FA: *p*_spin_ = 0.042 (females)).

**Figure S11.**
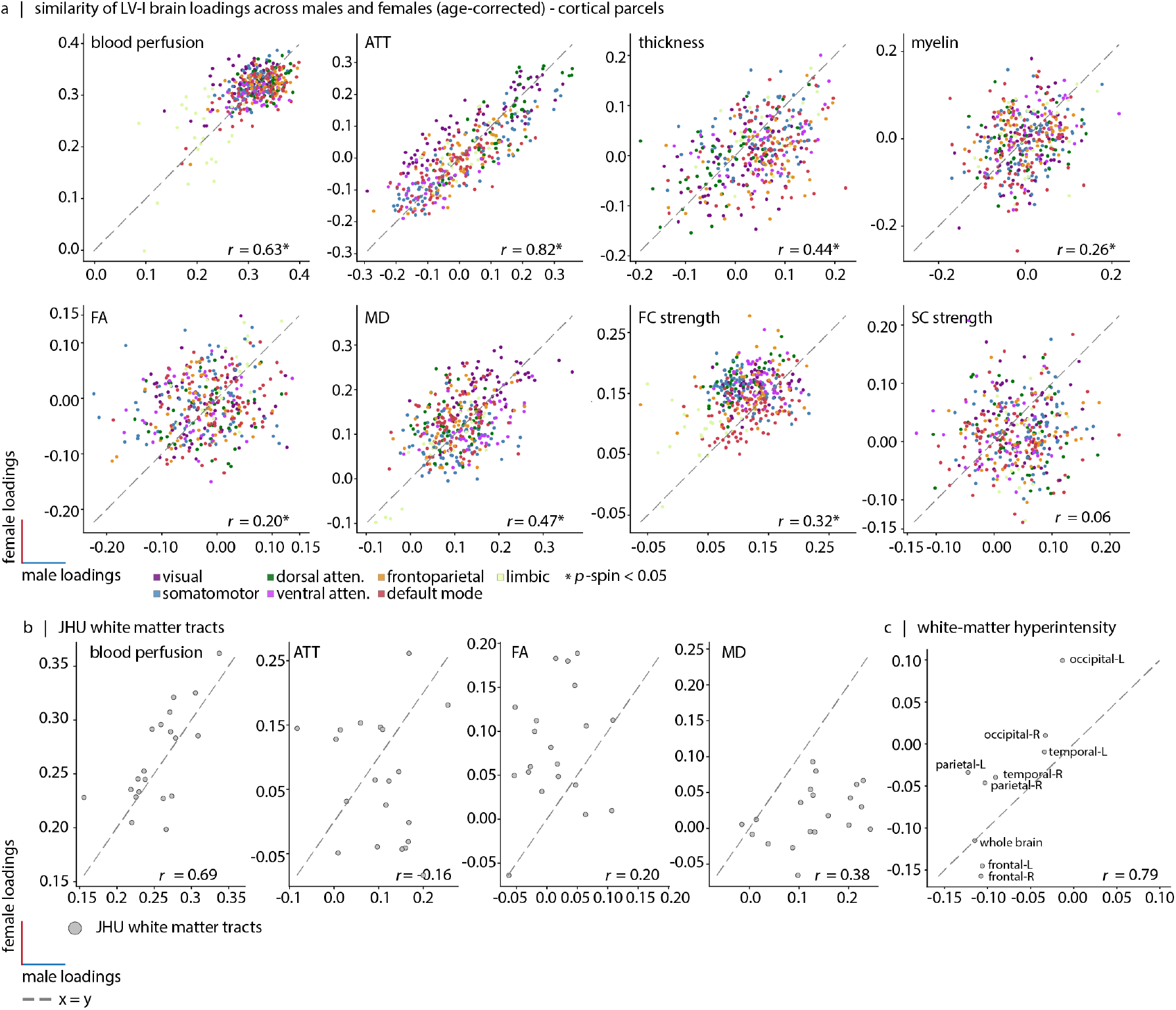
Similarity of first latent variable brain loadings between male and female - no age effect. (a) Cortical brain loadings comparison. In the scatter plots, each dot represents a brain region defined by the Schaefer-400 parcellation [182]; the dots are color-coded based on the Yeo-7 functional resting-state networks [208]. The *x*-axis shows male brain loadings and the *y*-axis shows female brain loadings. (b) White matter tract loadings comparison. The scatter plots show the similarity of brain loadings between males (*x*-axis) and females (*y*-axis) for white matter tract measurements including blood perfusion, arterial transit time (ATT), fractional anisotropy (FA), and mean diffusivity (MD). (c) White matter hyperintensity loadings comparison. The scatter plots show the similarity of brain loadings between males (*x*-axis) and females (*y*-axis) for white matter hyperintensity measurements.

**Figure S12.**
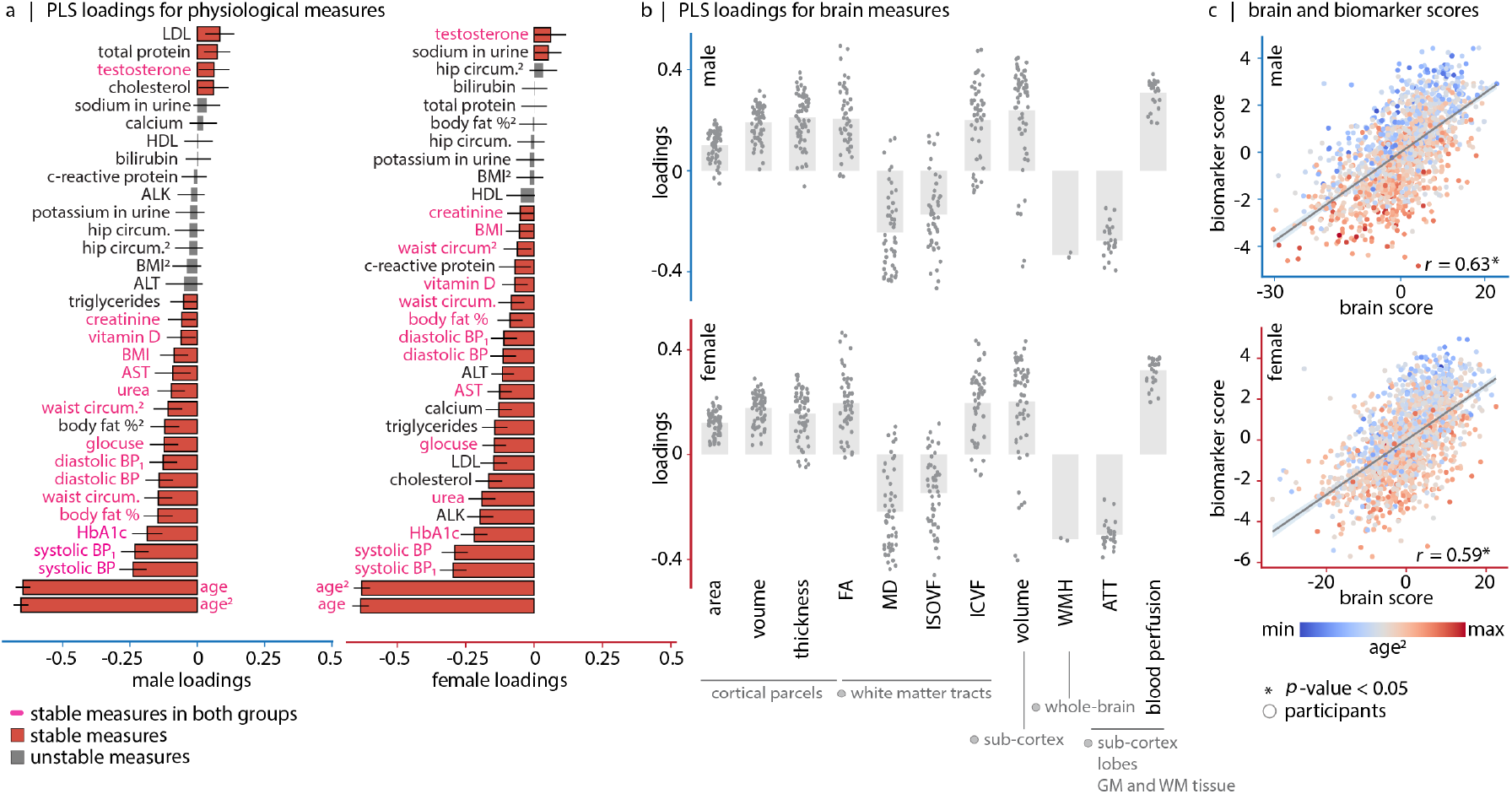
Mapping biomarkers to brain features in the UK Biobank dataset: first latent variable (LV–I) captures the aging axis. Using PLS analysis, we identify a significant latent variable that accounts for 75.71% (males) and 78.11% (females) of the covariance in the data. (a) Biomarker loadings. Bootstrap resampling is used to estimate the stability of each individual biomarker’s contribution to the overall multivariate pattern. Each biomarker loading is divided by its bootstrap-estimated standard error, yielding a measure called “bootstrap ratio”. Bootstrap ratio is high for biomarkers with large weights and small standard errors. Stable biomarkers for which the estimated 95% confidence interval do not cross zero, are shown in red. BMI, body fat percentage, age, hip and waist circumference were collected at both initial assessment and imaging visits. Variables measured in the imaging visit are denoted with a subscript _2_ (e.g., BMI_2_), while baseline values are written without subscripts. Superscript ^1^ is used to indicate repeated measurements (e.g., systolic blood pressure^1^). (b) Brain loadings. Each dot represents a brain region (cortical, subcortical, or a white matter tract). (c) Correlation between brain (*x*-axis) and biomarker scores (*y*-axis) for males (top; *r* = 0.63) and females (bottom; *r* = 0.59). The score per participant shows the extent to which the participant is expressing the brain-biomarker association captured by LV–I. Each dot represents an individual participant, colored by their age at imaging time-point. Score correlation values passed cross-validation in both sex groups.

**Figure S13.**
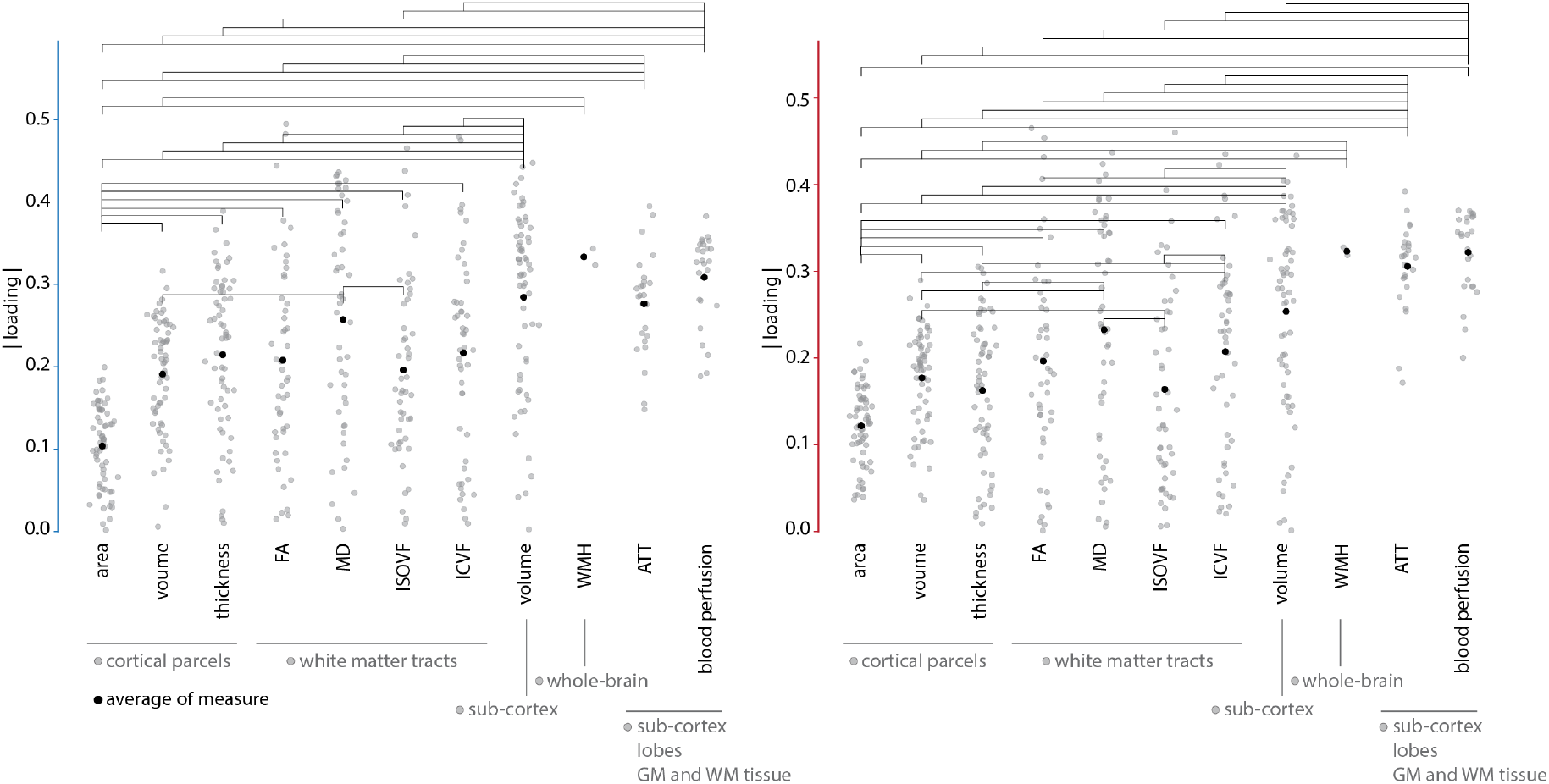
Statistical comparison of absolute brain loading values for the first latent variable in UK Biobank. Horizontal lines indicate statistical significance (*p <* 0.05, FDR corrected). Black dots represent mean absolute loading values. Also see Fig. S12b.

**Figure S14.**
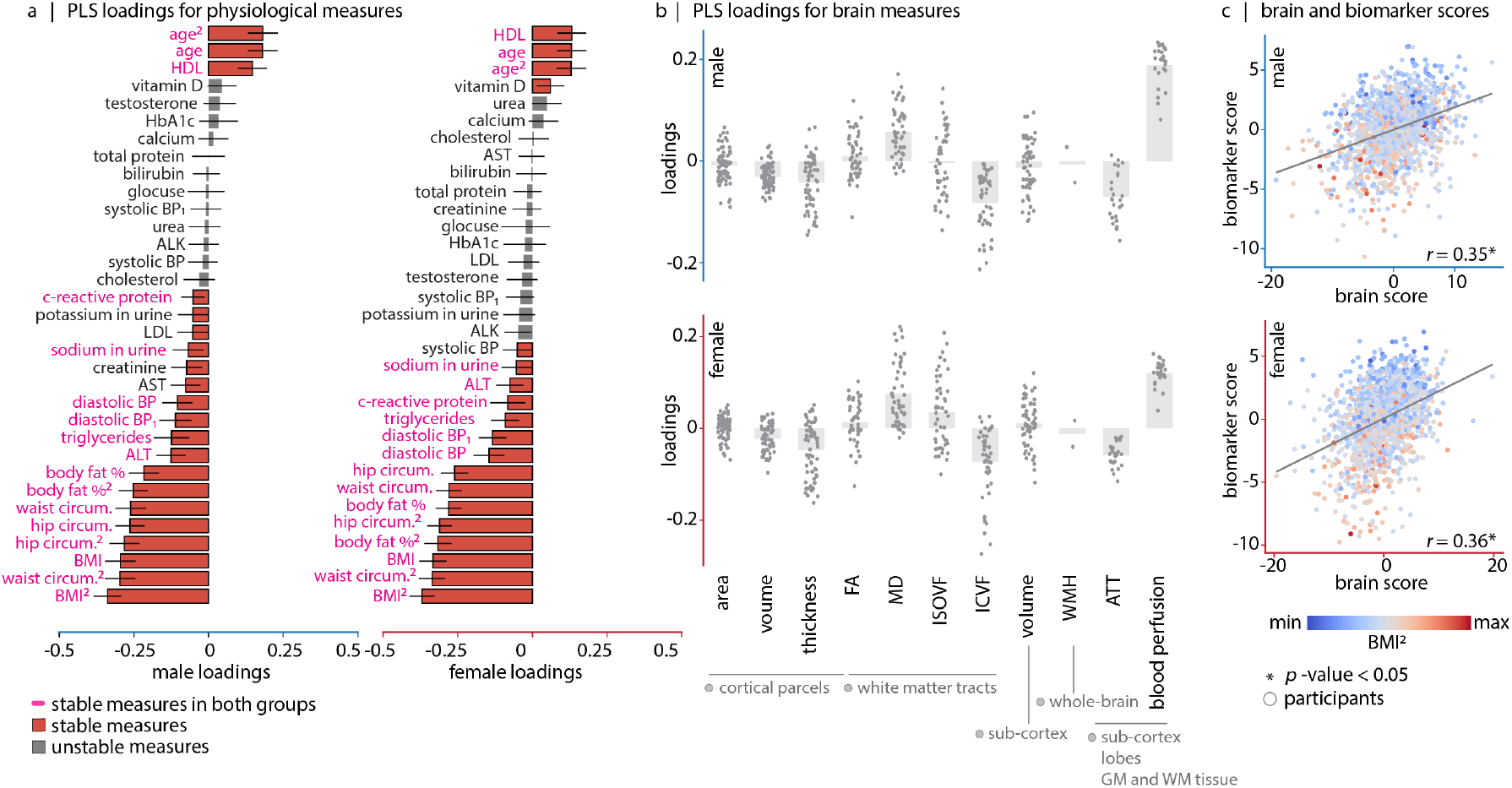
Mapping biomarkers to brain features in the UK Biobank dataset: second latent variable (LV–II) captures the metabolic axis. Using PLS analysis, we identify a significant latent variable that accounts for 15.33% (males) and 14.93% (females) of the covariance in the data. (a) Biomarker loadings. Bootstrap resampling is used to estimate the stability of each individual biomarker’s contribution to the overall multivariate pattern. Each biomarker loading is divided by its bootstrap-estimated standard error, yielding a measure called “bootstrap ratio”. Bootstrap ratio is high for biomarkers with large weights and small standard errors. Stable biomarkers for which the estimated 95% confidence interval do not cross zero, are shown in red. BMI, body fat percentage, age, hip and waist circumference were collected at both initial assessment and imaging visits. Variables measured in the imaging visit are denoted with a subscript _2_ (e.g., BMI_2_), while baseline values are written without subscripts. Superscript ^1^ is used to indicate repeated measurements (e.g., systolic blood pressure^1^). (b) Brain loadings. Each dot represents a brain region (cortical, subcortical, or a white matter tract). (c) Correlation between brain (*x*-axis) and biomarker scores (*y*-axis) for males (top; *r* = 0.35) and females (bottom; *r* = 0.36). Each dot represents an individual participant, colored by their BMI at imaging time-point. The scores per participant show the extent to which the participant is expressing the brain-biomarker association captured by LV–II. Score correlation values passed cross-validation in both sex groups.

**Figure S15.**
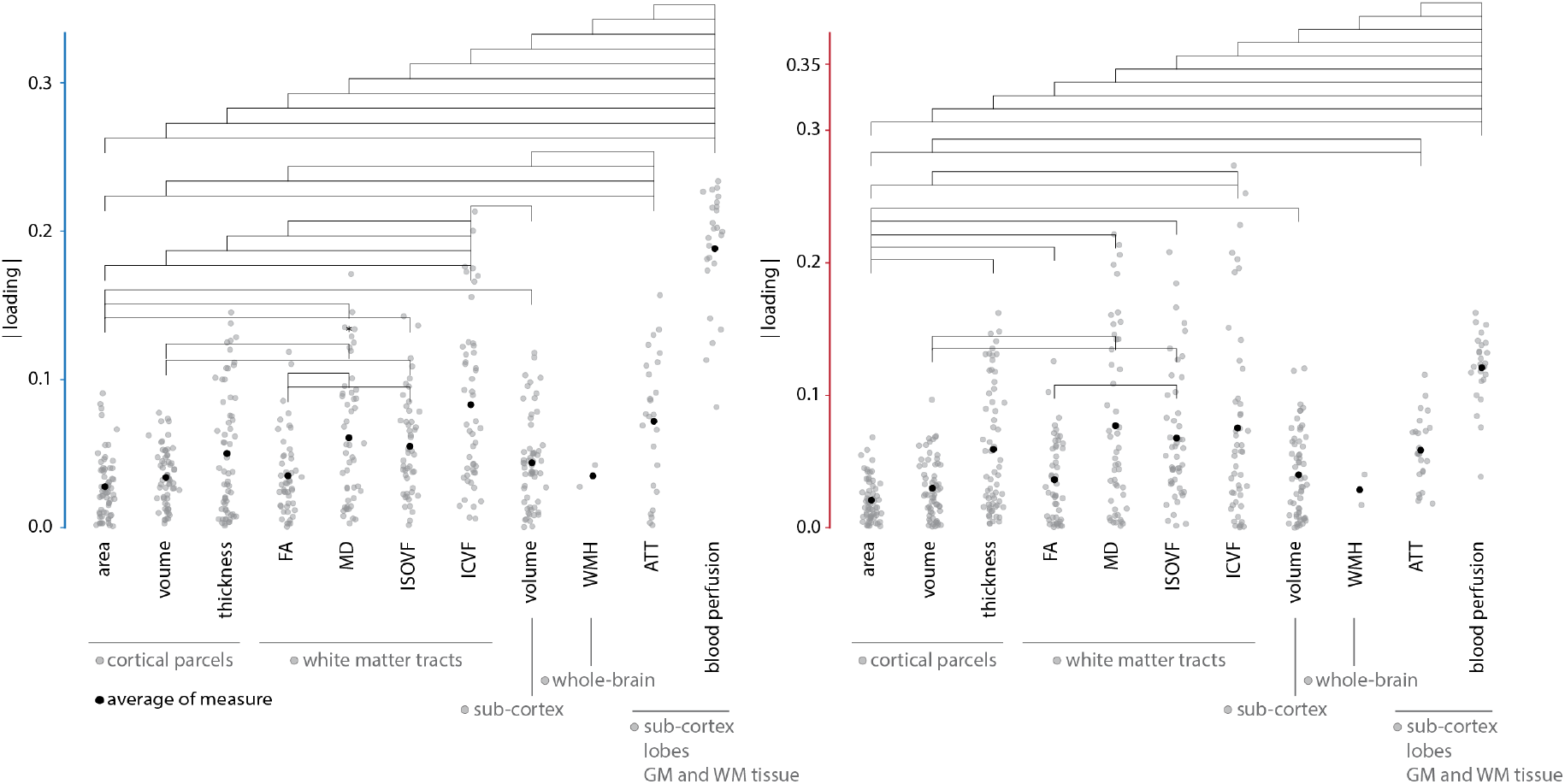
Statistical comparison of absolute brain loading values for the second latent variable in UK Biobank. Blood perfusion loadings have the greatest absolute values among all brain measures examined. Horizontal lines indicate statistical significance (*p <* 0.05, FDR corrected). Black dots represent mean absolute loading values. Also see Fig. S14b.

**Figure S16.**
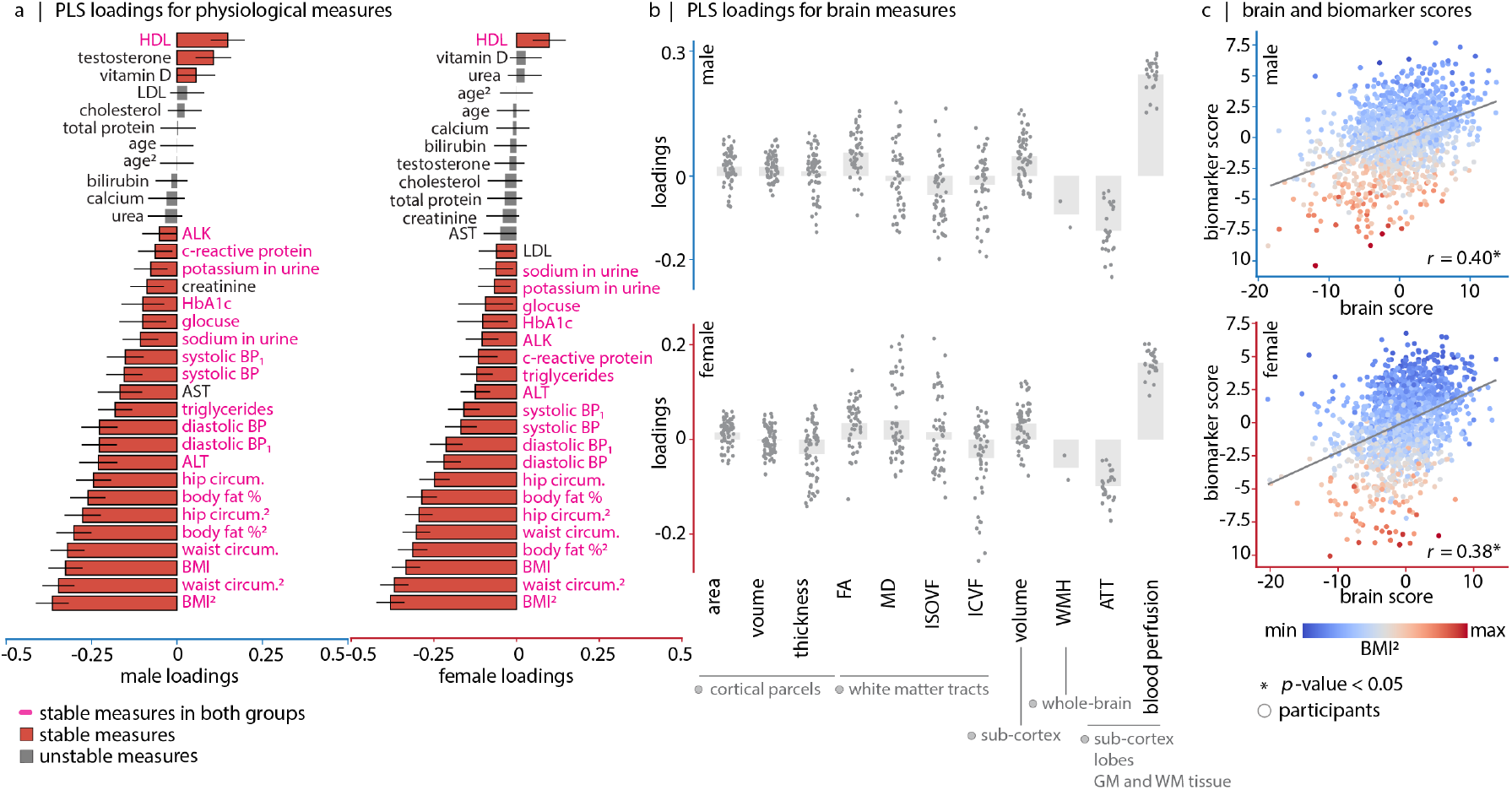
Mapping biomarkers to brain features after regressing out the age effect in the UK Biobank dataset: first latent variable (LV–I^*′*^) captures the aging axis. Using PLS analysis, we identify a significant latent variable that accounts for 65.31% (males) and 60.37% (females) of the covariance in the data. (a) Biomarker loadings. Bootstrap resampling is used to estimate the stability of each individual biomarker’s contribution to the overall multivariate pattern. Each biomarker loading is divided by its bootstrap-estimated standard error, yielding a measure called “bootstrap ratio”. Bootstrap ratio is high for biomarkers with large weights and small standard errors. Stable biomarkers for which the estimated 95% confidence interval do not cross zero, are shown in red. BMI, body fat percentage, age, hip and waist circumference were collected at both initial assessment and imaging visits. Variables measured in the imaging visit are denoted with a subscript _2_ (e.g., BMI_2_), while baseline values are written without subscripts. Superscript ^1^ is used to indicate repeated measurements (e.g., systolic blood pressure^1^). (b) Brain loadings. Each dot represents a brain region (cortical, subcortical, or a white matter tract). (c) Correlation between brain (*x*-axis) and biomarker scores (*y*-axis) for males (top; *r* = 0.40) and females (bottom; *r* = 0.38). Each dot represents an individual participant, colored by their BMI at imaging visit. The score per participant shows the extent to which the participant is expressing the brainbiomarker association captured by LV–I^*′*^. Score correlation values passed cross-validation in both sex groups.

**Figure S17.**
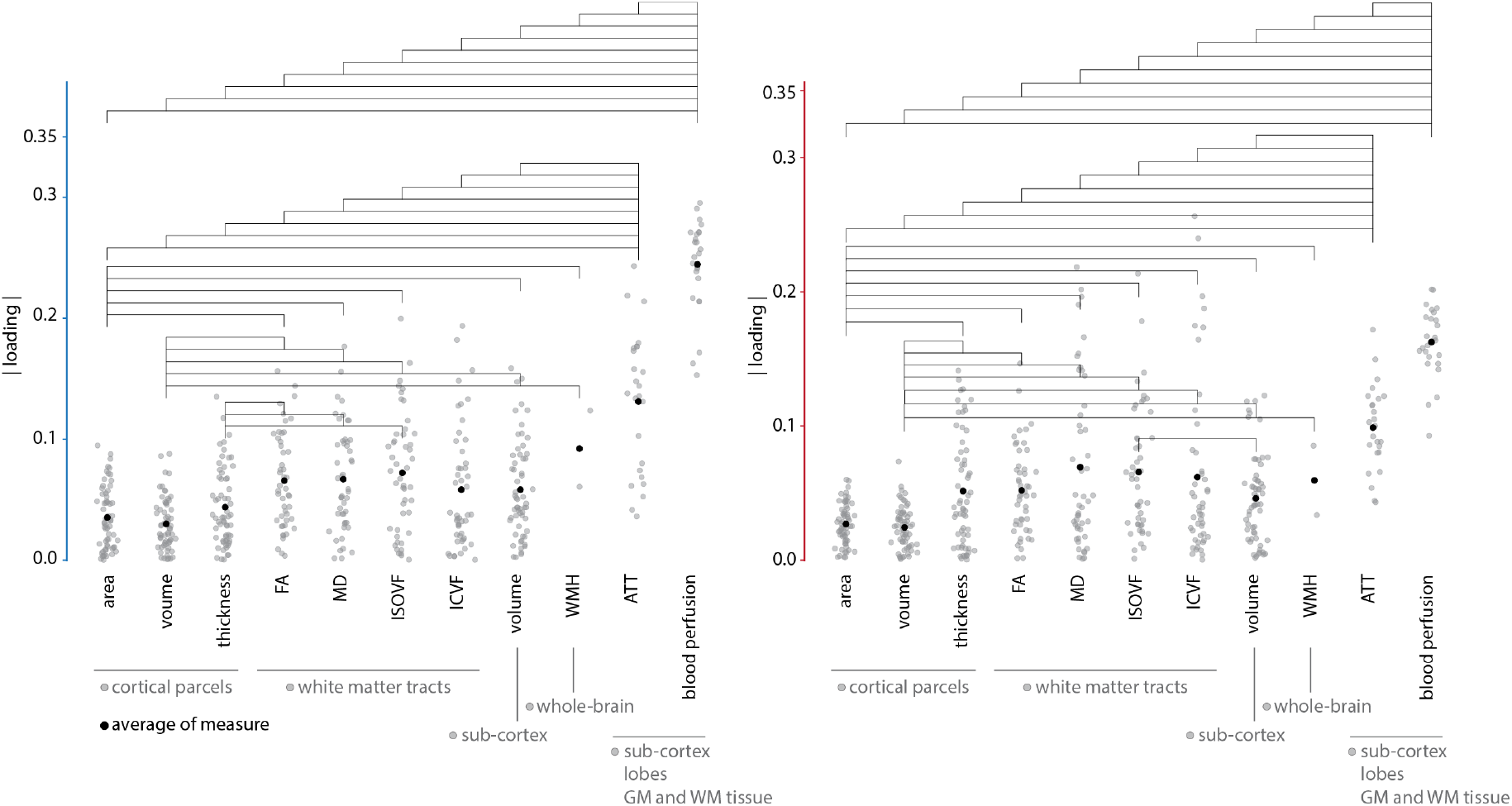
Statistical comparison of absolute brain loading values for the first latent variable in UK Biobank - no age effect. Blood perfusion loadings have the greatest absolute values among all brain measures examined. Horizontal lines indicate statistical significance (*p <* 0.05, FDR corrected). Black dots represent mean absolute loading values. Also see Fig. S16b.

**Figure S18.**
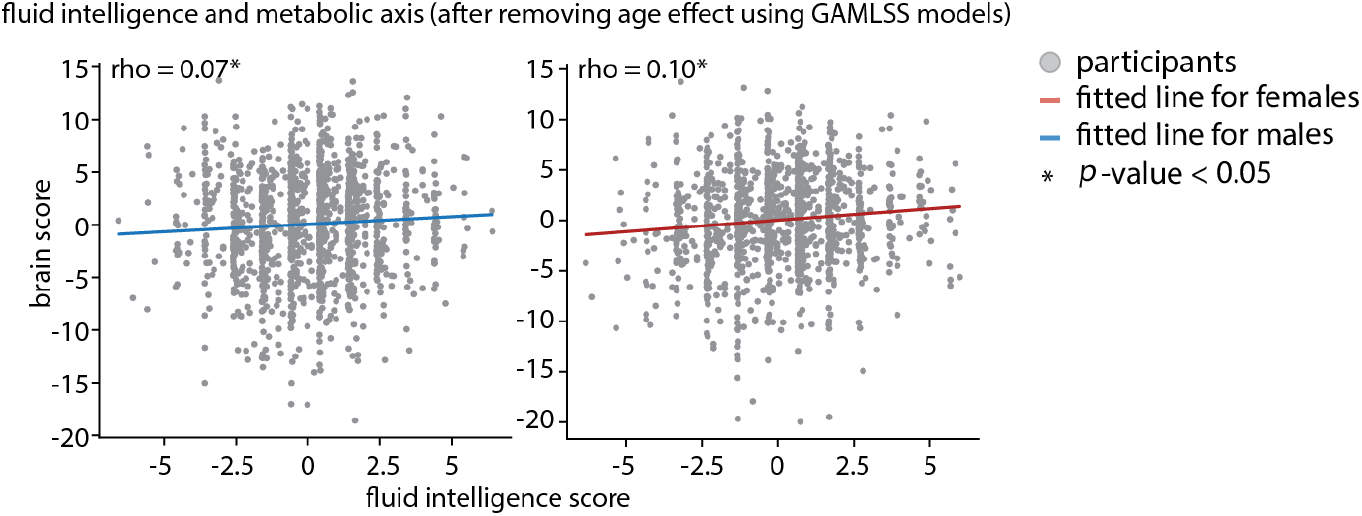
Cognitive relevance of metabolic axis in UK Biobank. The left scatter plot shows data for males (*p*_perm_ = 1.69 × 10^−2^) and the right scatter plot shows data for females (*p*_perm_ = 9.99 × 10^−3^) (*N*_perm_ = 1 000). Brain scores are derived from analysis presented in Fig. S16. Fitted regression lines are shown in blue for males and in red for females.

**TABLE S1.** Biomarkers in HCP–Aging. For each biomarker, the table reports the number of participants with missing values (NaN). Total number of females is 329, and total number of males is 268.

| **Biomarker** | **Number of NaN - female** | **Number of NaN - male** |
| --- | --- | --- |
| Age | 0 | 0 |
| Alanine aminotransferase (ALT) | 0 | 0 |
| Albumin | 0 | 0 |
| Alkaline phosphatase (ALK) | 0 | 0 |
| Aspartate aminotransferase (AST) | 0 | 0 |
| Bilirubin | 0 | 0 |
| Body mass index (BMI) | 2 | 0 |
| Calcium | 0 | 0 |
| Chloride | 0 | 0 |
| Creatinine | 0 | 0 |
| CO2 content | 0 | 0 |
| Estradiol | 1 | 2 |
| FSH | 1 | 2 |
| Glucose | 0 | 0 |
| Glycosylated haemoglobin (HbA1c) | 3 | 1 |
| HDL | 0 | 0 |
| Insulin | 0 | 0 |
| LDL | 0 | 0 |
| LH | 1 | 2 |
| Mean arterial pressure (MAP) | 4 | 2 |
| Potassium | 0 | 0 |
| Sodium | 0 | 0 |
| Testosterone | 1 | 2 |
| Total cholesterol | 0 | 0 |
| Total protein | 0 | 0 |
| Triglycerides | 0 | 0 |
| Urea | 0 | 0 |
| Vitamin D | 0 | 0 |

**TABLE S2.** Biomarkers in UK Biobank. For each biomarker, the table reports the acquisition session and the number of participants with missing values (NaN). Body mass index, body fat percentage, age, hip and waist circumference were measured at both the initial assessment and the imaging visits. Variables acquired during the imaging visit are denoted as 2.0 in the “Time point” column, while values acquired during the initial assessment are denoted as 0.0. A value of 0.1 indicates values acquired during the repeated assessment visit.

| Biomarker | Time point - instance | Number of NaN - female | Number of NaN - male |
| --- | --- | --- | --- |
| Age | 0.0 | 0 | 0 |
| Age | 2.0 | 0 | 0 |
| Alanine aminotransferase (ALT) | 0.0 | 131 | 105 |
| Alkaline phosphatase (ALK) | 0.0 | 130 | 103 |
| Aspartate aminotransferase (AST) | 0.0 | 136 | 108 |
| Bilirubin | 0.0 | 135 | 110 |
| Body fat percentage | 0.0 | 22 | 8 |
| Body fat percentage | 2.0 | 42 | 55 |
| Body mass index (BMI) | 0.0 | 4 | 2 |
| Body mass index (BMI) | 2.0 | 7 | 9 |
| C-reactive protein | 0.0 | 136 | 105 |
| Calcium | 0.0 | 255 | 105 |
| Cholesterol | 0.0 | 131 | 103 |
| Creatinine | 0.0 | 132 | 105 |
| Diastolic blood pressure | 0.0 | 128 | 115 |
| Diastolic blood pressure | 0.1 | 154 | 129 |
| Glucose | 0.0 | 257 | 197 |
| Glycosylated haemoglobin (HbA1c) | 0.0 | 143 | 116 |
| HDL | 0.0 | 254 | 195 |
| Hip circumference | 0.0 | 2 | 1 |
| Hip circumference | 2.0 | 3 | 7 |
| LDL | 0.0 | 133 | 106 |
| Potassium in urine | 0.0 | 67 | 46 |
| Sodium in urine | 0.0 | 65 | 44 |
| Systolic blood pressure | 0.0 | 128 | 115 |
| Systolic blood pressure | 0.1 | 154 | 129 |
| Testosterone | 0.0 | 325 | 117 |
| Triglycerides | 0.0 | 132 | 104 |
| Total protein | 0.0 | 255 | 195 |
| Urea | 0.0 | 131 | 104 |
| Vitamin D | 0.0 | 181 | 138 |
| Waist circumference | 0.0 | 2 | 1 |
| Waist circumference | 2.0 | 3 | 7 |

## Notes

https://github.com/netneurolab/Farahani_Metabolic_Health

### Competing Interest Statement

The authors have declared no competing interest.

### Summary of Updates

Revision Summary: The author information is updated.

